# Comparison of imaging-based single-cell resolution spatial transcriptomics profiling platforms using formalin-fixed, paraffin-embedded tumor samples

**DOI:** 10.1101/2024.12.13.628390

**Authors:** Nejla Ozirmak Lermi, Max Molina Ayala, Sharia Hernandez, Wei Lu, Khaja Khan, Alejandra Serrano, Idania Lubo, Leticia Hamana, Katarzyna Tomczak, Sean Barnes, Jinzhuang Dou, Qingnan Liang, RTI Team, Maria Gabriela Raso, Ximing Tang, Mei Jiang, Beatriz Sanchez-Espiridion, Annikka Weissferdt, John Heymach, Jianjun Zhang, Boris Sepesi, Tina Cascone, Anne Tsao, Mehmet Altan, Reza Mehran, Don Gibbons, Ignacio Wistuba, Cara Haymaker, Ken Chen, Luisa M. Solis Soto

## Abstract

Imaging-based spatial transcriptomics (ST) is evolving rapidly as a pivotal technology in studying the biology of tumors and their associated microenvironments. However, the strengths of the commercially available ST platforms in studying spatial biology have not been systematically evaluated using rigorously controlled experiments. In this study, we used serial 5-m sections of formalin-fixed, paraffin-embedded surgically resected lung adenocarcinoma and pleural mesothelioma tumor samples in tissue microarrays to compare the performance of the single cell ST platforms CosMx, MERFISH, and Xenium (uni/multi-modal) platforms in reference to bulk RNA sequencing, multiplex immunofluorescence, GeoMx Digital Spatial Profiler, and hematoxylin and eosin staining data for the same samples. In addition to objective assessment of automatic cell segmentation and phenotyping, we performed pixel-resolution manual evaluation of phenotyping to carry out pathologically meaningful comparison between ST platforms. Our study detailed the intricate differences between the ST platforms, revealed the importance of parameters such as tissue age and probe design in determining the data quality, and suggested reliable workflows for accurate spatial profiling and molecular discovery.

---

Studying the whole panorama of cellular and molecular interactions in tissues accurately and within their functional context is vital for understanding of health and disease. Spatial transcriptomics (ST) assays characterize gene expression profiles and localize them on histological tissue sections, preserving the context of interactions present in the tissue. *Nature Methods* selected ST as its Method of the Year 2020 [1], and this method has evolved rapidly since, with many technology companies including it in their assays [2]. ST has special significance in studying the association between a tumor and its microenvironment in cancer biology [3]. This novel technology can detect gene expression while preserving the location of genes at the single-cell level when using imaging-based ST techniques that rely on fluorescence in situ hybridization [4]. This allows researchers to investigate tissue sections and gain an understanding of complex interactions between cell populations and their arrangements within tissues [5].

Multiple commercial ST solutions have become available recently, such as CosMx Spatial Molecular Imaging (CosMx; NanoString, a Bruker company), MERFISH (Vizgen), and Xenium (10x Genomics), which are used to perform multiple cycles of nucleic acid hybridization of fluorescent molecular barcodes to identify RNA molecules while mapping their locations. However, they differ in their sample preparation protocols during amplification, gene selection for panel design, and cell-segmentation processes [6–9]. Comparison of imaging-based ST assays by multiple teams using different types of tissue is ongoing [9–11]. These studies will help researchers select optimal single cell ST methods for their assays, which are vital to producing high-quality data. In addition, we included pathologists’ evaluations of phenotyping results with guidance of H&E and mIF data of samples, a necessary step in understanding tissue morphology and determining the accuracy of these ST platforms in cell segmentation and cell-type annotation [12].

Herein, we compared the commercially available imaging-based ST platforms CosMx, MERFISH, and Xenium with unimodal (Xenium-UM) and multimodal (Xenium-MM) segmentation using 5-m serial sections of formalin-fixed, paraffin-embedded (FFPE) surgically resected lung adenocarcinoma and pleural mesothelioma samples obtained from 2016 to 2022 and placed in tissue microarrays (TMAs). We evaluated each platform by comparing multiple metrics, including tissue age, average transcript count and uniquely expressed gene count per cell, and signal detection above background by using negative control probes as well as blank probes. Later, we investigated the performance of cell segmentation across the platforms by evaluating the presence of transcripts in cells and individual cell area sizes as well as co-expression of disjoint genes by measuring the level of joint detections from genes that are mutually exclusive among cell populations. We also measured the concordance of the imaging-based ST data across the ST platforms with data obtained using bulk RNA sequencing (RNA-seq) and the GeoMx Digital Spatial Profiler with the Whole Transcriptome Atlas (DSP WTA) for the same cohort used with ST. In addition, we compared cell-type annotations among platforms based on selected genes in the panels of each ST platforms and pathologists’ evaluation of phenotyping against multiplex immunofluorescence (mIF) and hematoxylin and eosin (H&E) stained sections of samples.

The objectives of this study were to provide a comprehensive explanation of the differences in imaging, multiplexing and tissue age capability, and data acquisition among the three ST platforms using a platform-agnostic data analysis pipeline that includes dataset validation with single-cell RNA (scRNA)-seq data and a pathology-oriented review of the final cell type annotations produced by each platform’s probe reads. All comparisons were performed in a cutting-edge, high-throughput, high-complexity cancer research setting.

## Results

### Selection of lung adenocarcinoma (immune hot) and pleural mesothelioma (immune cold) data sets for comparison of the ST platforms

To ensure that our comparison of the performance of single-cell ST platforms can be used for translational studies of immuno-oncology research, we used archived FFPE tumor samples, which represent the current standard for sample processing and archiving in pathology. Specifically, we used four TMAs containing samples of surgically resected tumor tissue obtained in different years and with different immune profiling features: two TMAs containing samples of lung adenocarcinoma (ICON1 and ICON2; collected from 2016 to 2018), which is regarded as an immune hot tumor, and two TMAs containing samples of pleural mesothelioma (MESO1 and MESO2; collected from 2020 to 2022), which is regarded as an immune cold tumor. We submitted serial sections of each TMA to the ST companies to run the single-cell imaging–based ST assays. We subjected the ICON2 and MESO2 TMAs to all three ST assays, while ICON1 and MESO1 were unavailable in Xenium and MERFISH due to assay and tissue constraints, respectively. We used the best available panel for immuno-oncology research, which included genes relevant to lung adenocarcinoma and pleural mesothelioma (Fig. 1a).

**Fig. 1.**
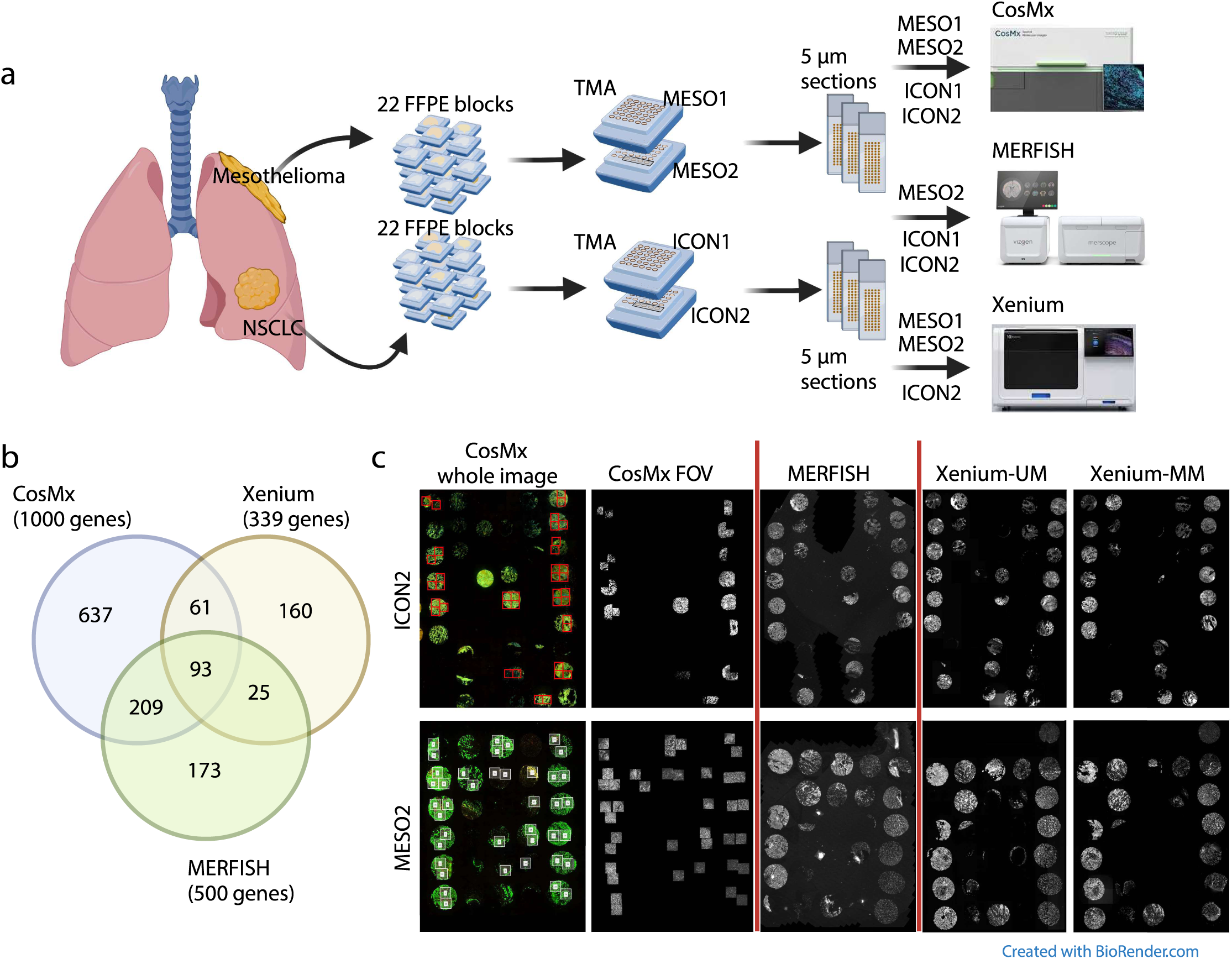
Experimental design, panel comparison, and nuclear staining with the ST platforms. **a,** TMAs containing tissue samples from patients with pleural mesothelioma (*n* = 22) or non-small cell lung cancer (NSCLC; *n* = 22). Tumor samples were sectioned at a thickness of 5 m and submitted to CosMx, MERFISH, Xenium-UM, and Xenium-MM assays based on tissue availability. **b,** Venn diagram of the genes shared by the panels of the ST platforms. **c,** CosMx whole images of the ICON2 and MESO2 TMAs stained for morphology markers before FOV selection. Red/white squares display selected FOVs of CosMx. The CosMx FOV, MERFISH, Xenium-UM, and Xenium-MM images display nuclear staining of selected FOV regions and whole tissue cores. Only TMAs with data available from all ST platforms were selected for comparisons.

### Comparison of gene panel coverage and imaging

The panels used in our study were the standard CosMx Human Universal Cell Characterization Panel (RNA, 1,000-plex), human MERSCOPE Immuno-Oncology Panel for MERFISH (RNA, 500-plex), and a 289-plex human lung panel plus a panel of 50 custom genes for Xenium, with 93 genes shared by all of the panels. The CosMx and Xenium panels shared 154 genes, the CosMx and MERFISH panels shared 302 genes, and the Xenium and MERFISH panels shared 118 genes (Fig. 1b and Supplementary Table 1).

The CosMx pipeline required region selection (field of view [FOV]) with 545 x 545-m squares, and we acquired up to 47 FOVs per slide in our experiment. Thus, we could not analyze whole tissue cores in CosMx data. The MERFISH and Xenium pipelines covered the whole tissue area mounted on each slide according to the manufacturers’ instructions (see Methods section) (Fig. 1c).

### Variation of transcript counts and unique gene detections per cell among the ST platforms

Transcript counts per cell were crucial to annotating cell types in the downstream analysis. We filtered cells with fewer than 30 transcript counts and that were five times larger than the geometric mean of cell area sizes of all cells on the CosMx platform based on the recommendations of NanoString. For analysis of MERFISH and Xenium, we removed cells with fewer than 10 transcript counts to avoid cells without any transcript counts or with low transcript counts. After cell filtering, we calculated the number of transcripts per cell and unique gene counts per cell in each TMA. The more recently constructed MESO TMAs had higher numbers of transcripts and uniquely expressed genes per cell with CosMx and MERFISH than Xenium. CosMx detected the highest transcript counts and uniquely expressed gene counts per cell among all TMAs, whereas MERFISH detected lower transcript and uniquely expressed gene counts per cell in the ICON1 and ICON2 TMAs than in the newer MESO2 TMA. However, Xenium-MM and Xenium-UM detected similar transcript and uniquely expressed gene counts per cell irrespective of TMA age. When comparing the two Xenium segmentation modalities, Xenium-UM assay had higher transcript and gene counts per cell than did Xenium-MM assay (Fig. 2a, b).

**Fig. 2.**
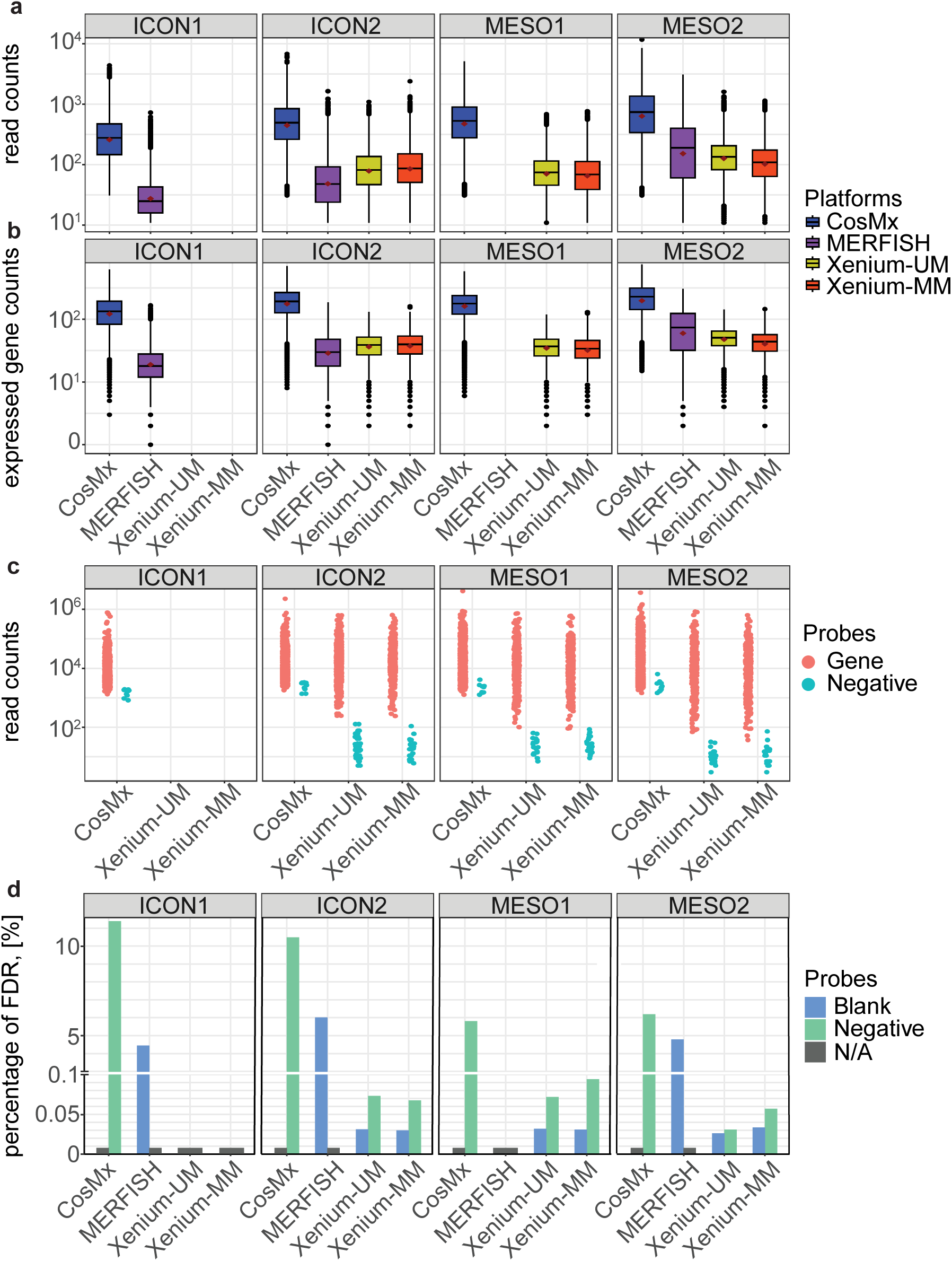
Technical comparison of the ST platforms. **a and b,** Box plots of the transcript counts and uniquely expressed genes per cell captured on each ST platform for each TMA tissue block. Each dot represents a cell. The red diamonds correspond to the average transcript counts and uniquely expressed genes in the blocks. **c,** Dot plot of the total transcript counts per probe across the ST platforms. Each dot represents a gene (red) or a negative control probe (green). **d,** Bar graph of FDRs calculated using negative or blank control probes and total read counts. The y-axis represents FDR as a percentage. The gray bars represent unavailable data. The blue and green bars represent FDRs calculated using blank and negative control probes, respectively.

### Comparison of the expression of negative control and gene probes

The CosMx panel included 10 negative control probes, the MERFISH panel included 50 blank probes, and the Xenium panel included 20 negative control probes, 41 negative control code words, and 141 blank code words. We compared the expression levels for the negative control probes with those for the target gene probes to determine whether the three ST platforms had highly expressed negative controls. CosMx displayed a significant overlap of the negative and target gene probes (ICON1, *n* = 8 [0.8%]; ICON2, *n* = 27 [2.7%]; MESO1, *n* = 196 [19.6%]; MESO2, *n* = 319 [31.9%]). Xenium-MM exhibited few overlapping genes (MESO2, *n* = 2 [0.6%]), whereas Xenium-UM did not have any genes overlapping the negative controls. Multiple overlapping probes in the CosMx dataset were important for cell-type annotation (e.g., *CD3D*, *CD40LG*, *FOXP3*, *MS4A1*, *MYH11*) (Fig. 2c, Supplementary Table 2). We could not perform this comparison using MERFISH owing to a lack of negative control probes.

We then assessed the specificity of each platform by calculating false discovery rates (FDRs) using negative control and blank code words/probes individually [9, 14–16]. We observed that CosMx (ICON, FDR > 10%; MESO, FDR > 5%) and MERFISH (ICON, FDR > 4%; MESO, FDR > 4%) had higher FDRs than did Xenium (both UM and MM: ICON, FDR < 0.07%; MESO, FDR < 0.09%). In addition, we observed similar FDRs in all Xenium TMAs (Fig. 2d).

### Impact of cell segmentation on average cell area size

Cell area sizes were crucial for interpreting transcript counts per cell. CosMx, MERFISH, and Xenium-MM included staining for morphological markers that are effective at identifying cell boundaries, and we processed ST data using standard cell-segmentation pipelines provided by the manufacturers. We performed cell segmentation with Xenium-UM via nuclear detection with a nuclear stain and cell boundary expansion algorithm. For this dataset, we tested different DAPI pixel-intensity settings to adjust the nuclear detection (50, 75, and 100 photoelectrons) and adjusted the maximum distance for cell membrane expansion from 7 m to 15 m to find the optimum cell-segmentation parameters with the Xenium Ranger software tool [17]. We accepted a Xenium-UM setting of 75-pixel intensity from the DAPI stain for nuclear detection and expansion distance of 7 m as the best parameters for cell segmentation.

Next, we calculated the average and median cell area size per cell on each ST platform. Even with our best assessment of the Xenium-UM segmentation parameters, it generated larger cell area sizes than did the cell-segmentation algorithms with morphology marker guidance in CosMx, MERFISH, and Xenium-MM. Xenium-MM produced the smallest average cell area size across all TMAs and the closest cell shapes in biology that we observed microscopically (e.g., elongated shapes of fibroblasts). The variation in cell area size was similar in the CosMx and Xenium data but was tight in the MERFISH data (Fig. 3a, b).

**Fig. 3.**
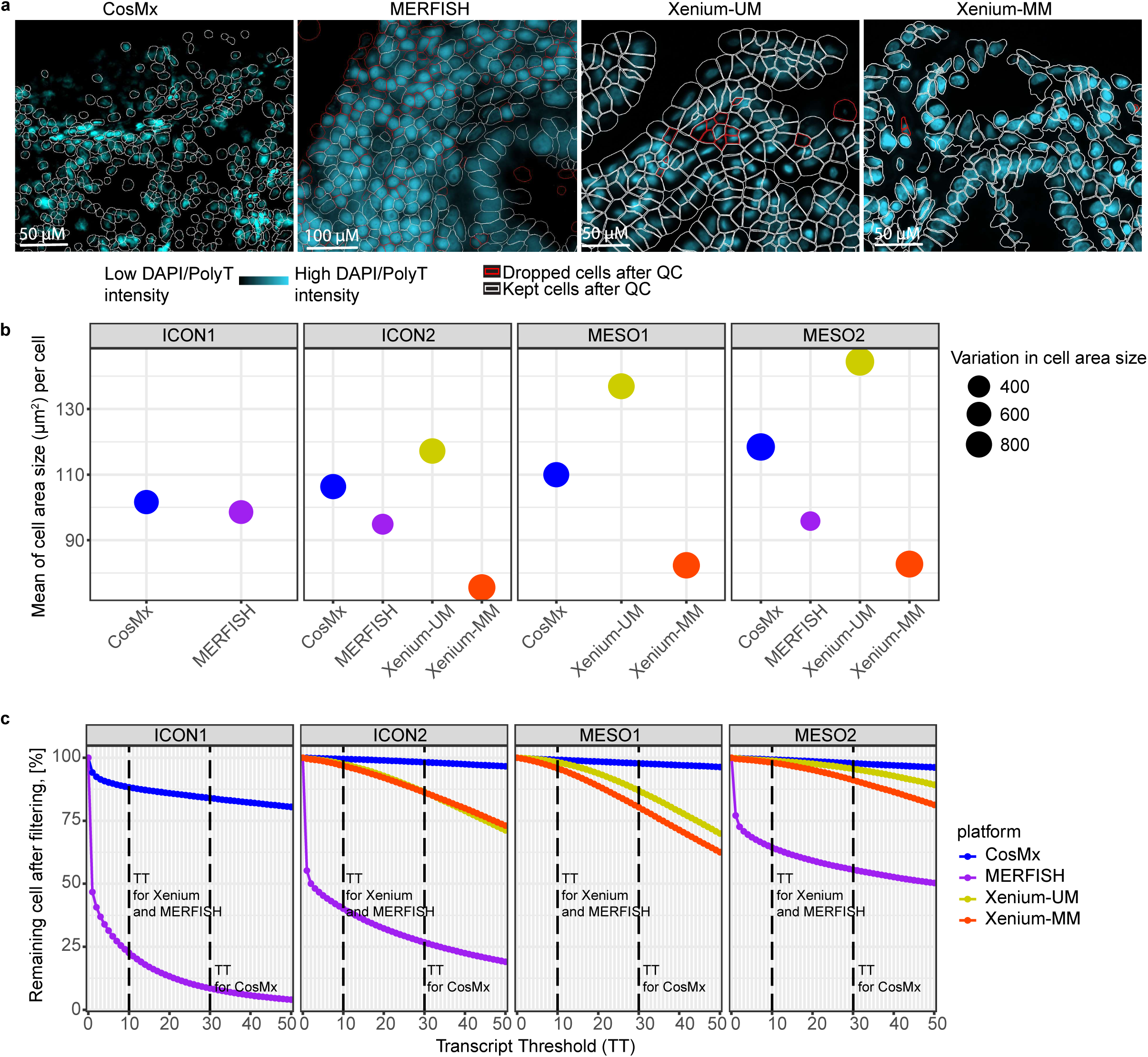
Comparative analysis of cell segmentation across the ST platforms. **a,** Images of nuclear staining overlaid with cell boundaries obtained on all platforms using the ICON2 TMA. Cells with red boundaries represent filtered cells after quality control. **b,** Bubble plot of the mean cell area sizes (m²) across the tissue blocks. The bubble size corresponds to the difference between the smallest and largest cell areas within each block. **c,** Percentages of the remaining cells after filtering. The dashed lines indicate transcript count thresholds for: Xenium (10), MERFISH (10), and CosMx (30).

### Impact of cell detection and segmentation on transcripts per cell and phenotyping

The fractions of cells after filtering with CosMx (ICON1, 84.0%; ICON2, 98.2%; MESO1, 97.8%; MESO2, 97.8%), Xenium-UM (ICON2, 97.4%; MESO1, 98.2%; MESO2, 98.8%), and Xenium-MM (ICON2, 96.8%; MESO1, 95.7%; MESO2, 98.0%) were greater than those after filtering with MERFISH (ICON1, 22.6%; ICON2, 40.0%; MESO2, 64.4%). The cell-segmentation step in MERFISH generated multiple cells without transcript counts located between the tissue cores or inside the tissue cores in the TMAs. Transcript counts in the segmented cells in MERFISH were absent from multiple tissue cores in the older TMAs (ICON1 and ICON2). CosMx had more filtered cells in the oldest TMA (ICON1) than in the other TMAs (Fig. 3a, c).

Next, we chose canonical markers for distinct cell populations in pairs, such as the canonical B-cell marker *CD19* and canonical T-cell marker *CD3E*; *EPCAM*, a marker for epithelial cancer cells; *CD4* and *CD8A*, which are markers for T-cell subsets; and *CD68*, a canonical myeloid/macrophage marker. We combined all filtered cells across all TMAs on each ST platform and excluded cells that did not express any cell marker from each of the corresponding gene pairs. All three platforms had multiple cells that co-expressed gene sets, and the number of cells with this co-expression increased when the differences in biological cell area size between cell types increased (e.g., T cells and tumor cells, T cells and macrophages). Xenium-MM performed better than the other platforms with comparison of multiple gene pairs. MERFISH performed better than the other platforms with the *EPCAM*-*CD3E* and *CD68*-*CD3E* gene pairs, but the number of detected cells with MERFISH was lower than those with the other platforms (Supplementary Table 3).

### Correlation of gene expression results with bulk RNA-seq and ST whole-transcriptome assay

The correlation of imaging-based ST platforms data with bulk RNA-seq was positive among all data sets. Whereas the concordance of ICON2 was lower for CosMx (*R* = 0.43, *P* < 0.0001) and MERFISH (*R* = 0.39, *P* < 0.0001) than Xenium-MM, the same correlation in the newest TMA (MESO2) was higher for both CosMx (*R* = 0.58, *P* < 0.0001) and MERFISH (*R* = 0.64, *P* < 0.0001) than ICON2 Cox and MERFISH. On the other hand, the concordance of ICON2 for Xenium-UM (*R* = 0.62, *P* < 0.0001) and Xenium-MM (*R* = 0.62, *P* < 0.0001) was lower than the concordance in MESO2 for Xenium-UM (*R* = 0.66, *P* < 0.0001) and Xenium-MM (*R* = 0.65, *P* < 0.0001), but the difference between ICON2 and MESO2 correlation of Xenium-Um with bulk RNA-seq was not as big as what we observed for CosMx and MERFISH (Fig. 4a,b).

**Fig. 4.**
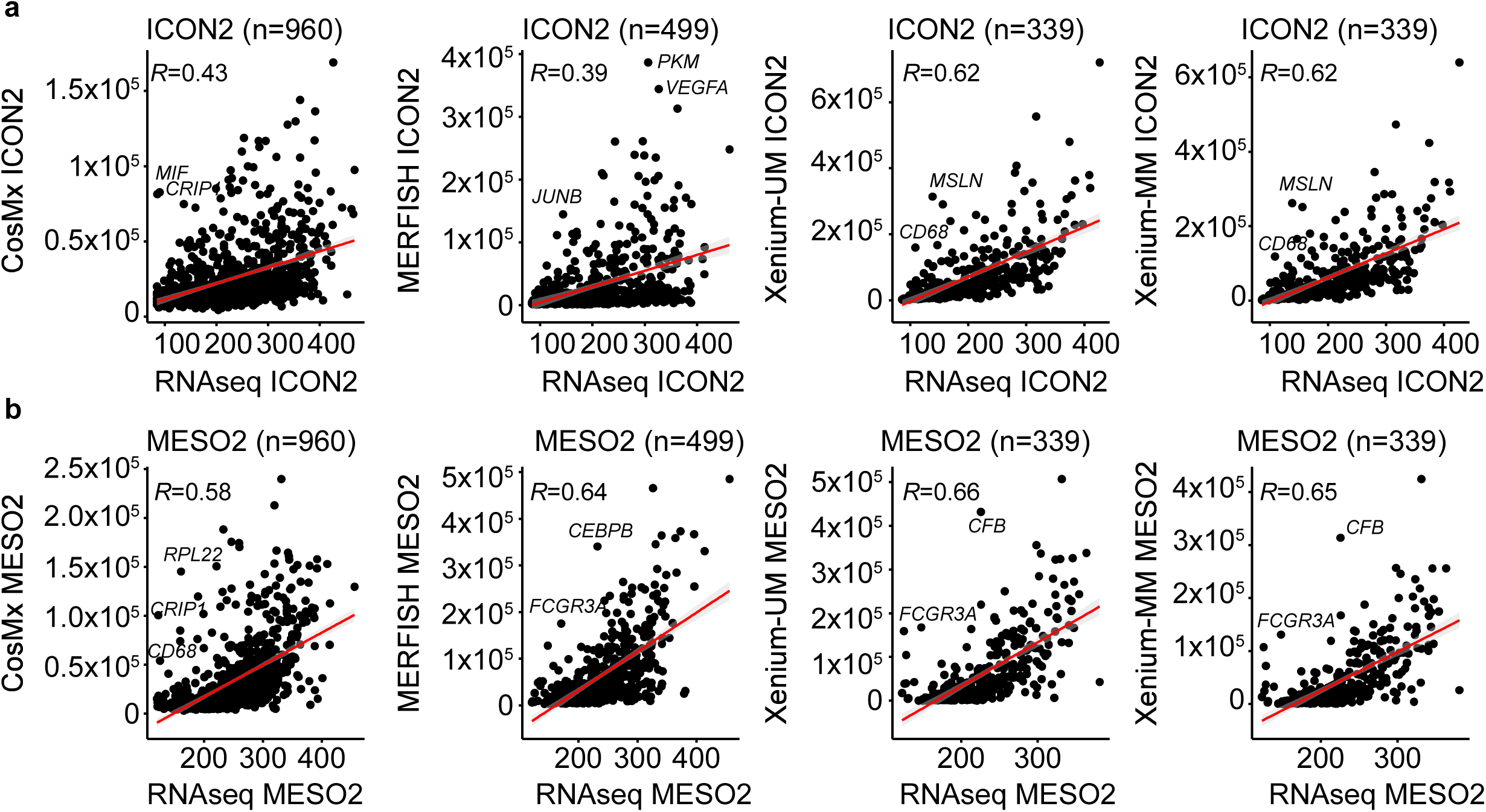
Concordance of RNA levels between ST platforms and bulk RNA-seq data. Scatter plots of the average expression of overlapping genes in ST platform and bulk RNA-seq data for matched cohorts of the ICON2 (**a**) and MESO2 (**b**) TMAs. The red lines represent linear regression. Pearson correlation coefficients (*R*) are provided in the top left corner of each plot. Outlier genes are annotated.

Afterward, we compared ST data with the GeoMx DSP WTA results. CosMx data (ICON2: *R* = 0.84, *P* < 0.0001; MESO2: *R* = 0.85, *P* < 0.0001) had better correlation with DSP WTA data than did MERFISH (ICON2: *R* = 0.67, *P* < 0.0001; MESO2: *R* = 0.77, *P* < 0.0001), Xenium-UM (ICON2: *R* = 0.70, *P* < 0.0001; MESO2: *R* = 0.80, *P* < 0.0001), and Xenium-MM data (ICON2: *R* = 0.68, *P* < 0.0001; MESO2: *R* = 0.80, *P* < 0.0001) (Extended Data Fig. 1).

We then compared the ST platforms with each other using shared genes to examine the reproducibility of ST platforms. CosMx and Xenium-UM data (ICON2: *R* = 0.86, *P* < 0.0001; MESO2: *R* = 0.88, *P* < 0.0001) correlated better than did CosMx and MERFISH (ICON2: *R* = 0.74, *P* < 0.0001; MESO2: *R* = 0.79, *P* < 0.0001) and Xenium-UM and MERFISH (ICON2: *R* = 0.67, *P* < 0.0001; MESO2: *R* = 0.84, *P* < 0.0001). Xenium-UM and Xenium-MM correlated strongly in ICON2 and MESO2 tissue blocks (*R* = 0.99, *P* < 0.0001). High correlation between Xenium-UM and Xenium-MM demonstrated the reproducibility of Xenium data even when using different cell-segmentation methods (Extended Data Fig. 2).

### Differences in clustering and cell-type annotations among ST platforms

To compare each ST platform’s ability to phenotype cells accurately, we used two different approaches for each set of TMAs. With the ICON1 and ICON2 TMAs, we used a manual clustering-based algorithm with a series of lineage markers that we selected based on the available genes in each panel. However, we excluded no markers from the panels even if target genes were within the range of expression of the negative control probes.

Using CosMx and the ICON TMAs, we were able to identify nine major cell types: macrophages, monocytes, fibroblasts, tumor cells, endothelial cells, smooth muscle cells, plasma cells, mast cells, and T cells. We labeled a cluster as being of unknown cell type if it had expression of *CD19*, *CD3E* and *EPCAM* in the same cluster. T-cell cluster had low expression of B-cell markers (e.g., *CD19*, *MS4A1*). Upon further investigation, we concluded that signals from *CD3E* and *CD19* came from every cell type in the clustering analysis, thus rendering these phenotypes undetectable. Moreover, correlation analysis of the assigned cell types of CosMx with ICON2 TMA demonstrated high levels of correlation between different cell types in multiple genes that distinguished each cluster (Fig. 5a and Extended Data Fig. 3).

**Fig. 5.**
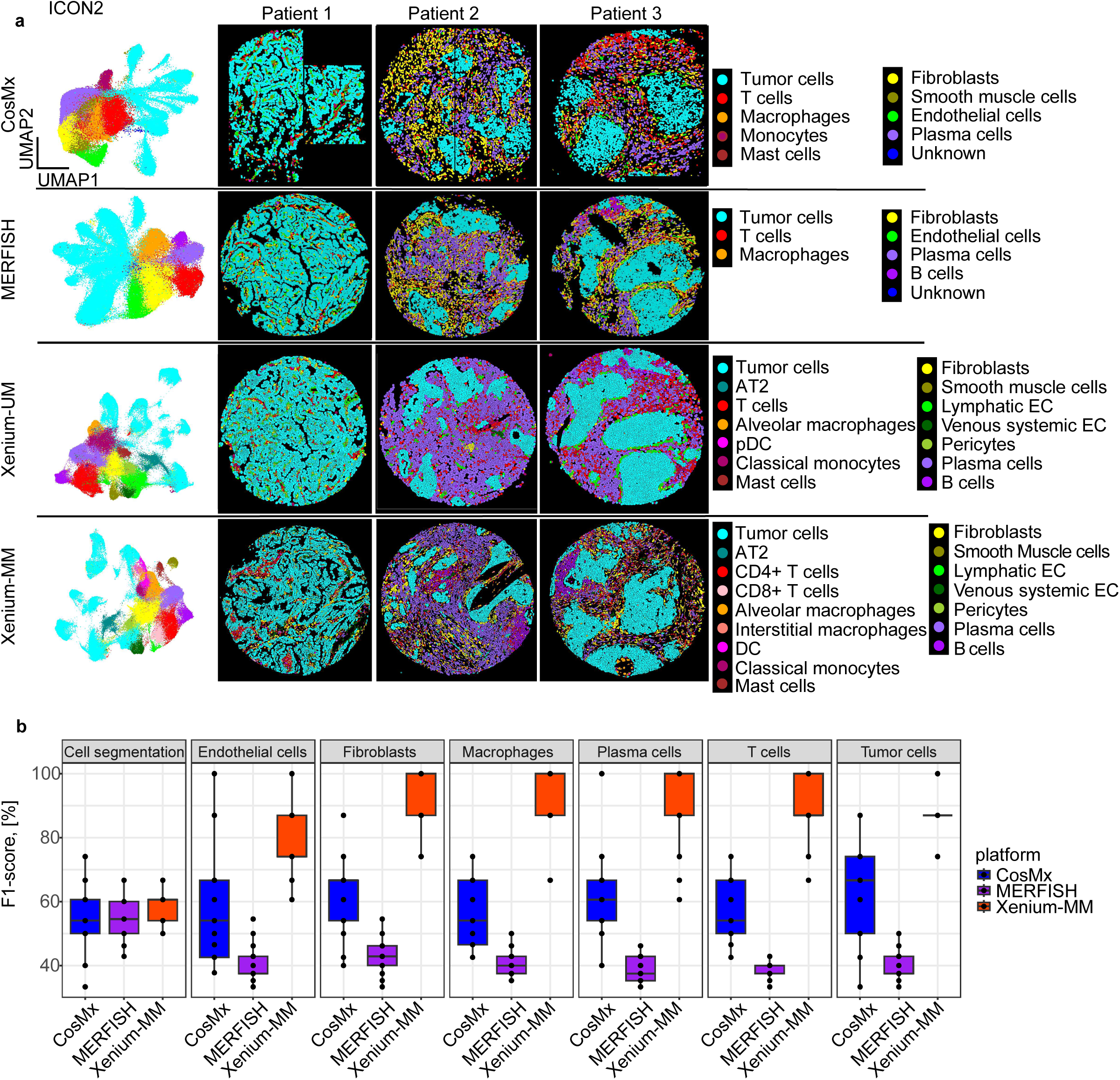
Cell-type annotation performance of the ST platforms using the ICON2 TMA. **a,** UMAP and spatial locations of manually annotated cell-type clusters across the platforms. Clusters are labeled with their corresponding cell types based on top-expressed genes. **b,** Box plot of the F1-scores representing performance of cell segmentation and cell type annotations in ST platforms. Each dot represents a tissue core.

For MERFISH, we excluded ICON1 data from our downstream data analysis owing to low transcript counts and missing cell segmentation in tissue cores and thus analyzed only ICON2 data. We detected six major cell types in MERFISH ICON2 data: macrophages, fibroblasts, endothelial cells, plasma cells, T-cells, and B-cells. We also identified an unknown cell type owing to low gene expression and a lack of differentially expressed genes to annotate the cluster using the manual clustering-based annotation method. Correlation analysis of the cell types displayed higher correlation among macrophages, fibroblasts, plasma cells, and T-cells than expected (Fig. 5a and Extended Data Fig. 4).

Using Xenium-UM with the ICON2 TMA, we detected 14 cell types via manual annotation: alveolar macrophages, classical monocytes, pericytes, alveolar type 2 cells, fibroblasts, plasmacytoid dendritic cells, lymphatic endothelial cells, venous system endothelial cells, smooth muscle cells, tumor cells, mast cells, plasma cells, T cells, and B cells. Differentially expressed genes in each cluster were helpful for annotation with a specific cell type. Xenium-UM was effective at differentiating some cell subtypes, such as lymphatic and venous system endothelial cells (Fig. 5a).

Regarding Xenium-MM with ICON2, we detected dendritic cells, T-cell subsets (CD4^+^ and CD8^+^), and macrophages (interstitial and alveolar) in addition to Xenium-UM via manual clustering-based annotation, but we did not detect plasmacytoid dendritic cells (Fig. 5a).

Correlation analysis of the different cell types demonstrated distinct separation between specific cell lineages. We annotated clusters of unique cell types even with the limited number of genes offered by the panel design of the ST platform (Extended Data Fig. 5, 6).

Next, we correlated assigned cell types on each platform in the ICON2 TMA using genes shared by the three ST platforms. Assigned cell types in the CosMx ICON2 data were strongly correlated with the same and other assigned cell types in the MERFISH and Xenium-UM ICON2 data. Assigned cell types in the MERFISH data were weakly correlated with the cell types in the Xenium-UM data. Examination of the correlations between assigned cell types in the Xenium-UM and Xenium-MM data demonstrated strong correlations between the same cell types and weak correlations between distant cell lineage types (Supplementary Fig. 1).

For our second approach, a label transfer method from scRNA-seq of 11 pleural mesothelioma samples and 5 adjacent normal pleural mesothelioma samples from the same cohort of MESO1 and MESO2 TMAs. We annotated some cells as unknown owing to low prediction scores (<10%). Although CosMx and MERFISH had overlapping gene expression between the multiple cell types, label transferring method from matched scRNA-seq performed better than manual cluster-based cell type annotation with pleural mesothelioma TMAs (Fig. 6 and Extended Data Fig. 7, 8). For MESO2 TMA, Xenium-UM and Xenium-MM were equally successful in annotating cell types by indicating the distinct separation among the cell types and lineages (Extended Data Fig. 9, 10).

**Fig. 6.**
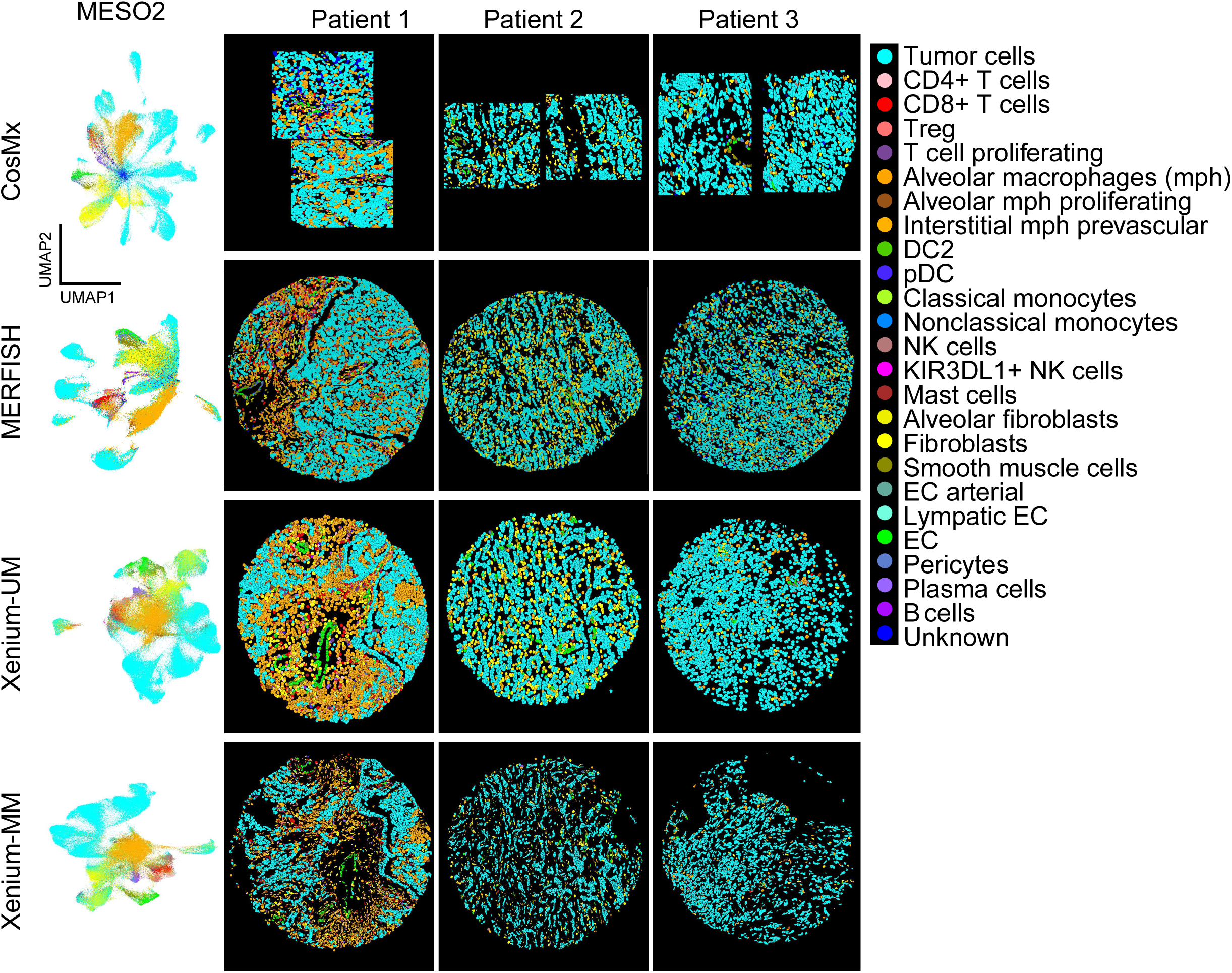
Cell-type annotation performance of the ST platforms in the MESO2 TMA. UMAP and spatial locations of annotated cell types across the platforms. The cell-type labels were transferred from scRNA-seq data on matched pleural mesothelioma cohorts using Seurat in R.

Finally, we examined the correlation of the assigned cell types on each ST platform in the MESO2 TMA using overlapping genes as we performed for the ICON2 TMA. Automated cell-type annotation using matched scRNA-seq of pleural mesothelioma samples produced better correlations of the same cell types on the platforms than CosMx and MERFISH. Xenium-UM and Xenium-MM displayed almost perfect correlations for the same cell types (Supplementary Fig. 2).

### Visual evaluation of cell phenotyping and segmentation

To assess the performance of cell phenotyping and segmentation on each ST platform, we compared the ST images with cell-type annotations produced by each platform for the ICON2 TMA with the images of the serial sections of TMAs stained with H&E and mIF for Syto13 (nuclei), CK (epithelium), CD3 (T cells), and CD68 (macrophages). Three independent pathologists experienced in immune profiling and image analysis assigned a score (range, 1-5; lower score represents greater similarity of histological/mIF features) to each tissue core for the overall cell segmentation and the following annotated cell types: epithelial cells, plasma cells, T cells, macrophages, endothelial cells, and fibroblasts. Next, we calculated the F1-score to measure of predictive performance of ST platforms (range, 0-1), accepting the average scores of pathologists’ evaluation as the ground truth, and compared the F1 results for all three platforms (Supplementary Fig. 3). The basis for this approach is that the location and morphology of these cell types in the images with cell-type annotations should have features comparable with those observed with H&E staining and/or mIF. We excluded Xenium-UM from pathologists’ evaluation due to lack of cell membrane detection for cell segmentation. Our results demonstrated that that F1-scores [18] for cell segmentation were similar among platforms. CosMx had higher variation in evaluation of cell segmentation, but in general, cell segmentation among the platforms was acceptable. However, Xenium-MM was the only platform with its exact section in H&E staining in the evaluation. We observed better F1-scores with Xenium-MM (median, >75%) than other ST platforms for all cell types; thus, Xenium-MM data had highly consistent standard morphological evaluation and cell-type annotation results. CosMx had the greatest range of F1-scores, demonstrating that the platform struggled to produce annotations that matched the morphology in certain scenarios. Additional exploration of these scenarios is needed to explain the factors that produce this wide range of agreement. MERFISH had the lowest F1-scores for all cell types (Fig. 5b).

## Discussion

In this study, we performed a systematic comparison of three commercially available single-cell imaging–based ST platforms using archived FFPE lung adenocarcinoma and pleural mesothelioma samples included in TMAs constructed from tissue blocks collected in different years and with annotated information of bulk RNA-seq and digital spatial profiling for matched tissue samples. The information on the tissue samples provided by these cutting-edge techniques highlights differences in panel size and gene coverage at the time of the evaluation, imaging workflows, and assay performance by including sensitivity and specificity (background) of probes, cell segmentation, and phenotyping information. Furthermore, we integrated histopathological evaluation of the patterns obtained from different cell phenotypes with expected cellular patterns. All of the features that we evaluated in this study are key considerations for experimental design of single-cell imaging–based ST assays for immuno-oncology research.

The ST platforms had multiple differences in performance. The first difference was in the ability to produce data on aged tissue samples. Whereas CosMx and MERFISH performed better with newly processed TMAs, Xenium performed uniformly across different tissue ages. The panel sizes and gene selections that we used differed substantially (CosMx, 1,000-plex; MERFISH, 500-plex; Xenium, 289-plex lung panel + 50 custom genes). Only 93 genes were shared by all three platforms. Researchers should investigate the panel design before running an assay to understand whether the panel covers their genes of interest. CosMx covered the most genes in the panel; hence, we observed the highest transcript counts and highest number of uniquely expressed genes per cell with CosMx.

Tissue preparation and coverage of whole tissue area was challenging. CosMx required less training to mount tissue samples to Superfrost slides than MERFISH and Xenium for mounting tissue to circular and Xenium-specific slides, respectively. After tissue samples were mounted to assay-specific slides, selection of the FOV under the guidance of a pathologist using morphology marker staining was required with CosMx. This step reduced the coverage area on this platform. In this study, we used TMAs constructed from multiple samples; thus, CosMx did not cover some of the tissue cores owing to the selection of FOVs. We analyzed whole tissue cores mounted to slides using MERFISH and Xenium.

Negative control and blank probes were necessary for estimating the background signal of the assays. We observed high expression of negative control probes in the CosMx assays. We could not include MERFISH in this comparison owing to a lack of negative control probes. High expression of negative controls represented the background or noise level in the CosMx assays. Later, we calculated FDRs on each platform using negative control and blank probes separately for each assay to estimate the error rates on each platform. The FDRs for CosMx and MERFISH were high in all TMAs. Xenium displayed low expression of negative controls in all TMAs; thus, the FDR for Xenium with both negative control and blank probes was low.

Accomplishing accurate cell segmentation has been a hurdle with single-cell imaging– based ST platforms. Each platform tries to solve the problem of cell segmentation using different strategies. We found that cell-segmentation algorithms with morphology marker staining performed better than the cell boundary expansion method did. However, these algorithms depended on the quality of morphology marker staining. When a problem in the staining step was encountered, cell segmentation was inconsistent owing to the background for the morphology marker staining and generated cells without transcript counts as we observed with the MERFISH assay. Xenium-UM was the only method without morphology marker staining, and it had inaccurate cell segmentation, especially in tissue regions with air spaces. Although we reduced the cell size for Xenium-UM to no more than 7 m in the cell segmentation step, the cells were larger than other assays with morphology marker staining. Inaccurate cell segmentation causes an incorrect assigning of transcripts to neighbor cells.

Precise cell-type annotation was crucial for downstream analysis and was affected by all previously mentioned steps above, including panel design and cell segmentation. We performed a cluster-based manual approach to annotate cell types with lung adenocarcinoma samples when matched scRNA-seq results were not available. We considered different cell markers to annotate cell types manually on each platform owing to differences in panel design. The noise level in the CosMx assay eliminated the distinct highly expressed genes in the clusters; thus, we could not separate B cells from T cells and labeled a cluster as unknown because of the expression of multiple distinct cell markers in CosMx. MERFISH had low gene expression; thus, one of the clusters in MERFISH did not have a clear cell-type indicator and was labeled as unknown. The low noise level in Xenium helped us to easily annotate clusters with a cell type based on available gene expression. Better cell segmentation with Xenium-MM than Xenium-UM increased the level of cell-type annotation from coarse cell lineages to cell subtypes for T cells. We tried to annotate lung adenocarcinoma samples with public reference datasets and an automated cell-type annotation algorithm using the TransferData() option in Seurat, but the results were inadequate owing to differences between cohorts of reference dataset and ST data. For the pleural mesothelioma samples, we benefited from having matched scRNA-seq data to annotate the ST data via Seurat co-embedding and label transfer. CosMx and MERFISH had low prediction scores for some cell types owing to high noise levels or low gene expression; hence, we labeled the cell types as unknown as we did previously. Having matched scRNA-seq data increased the accuracy of cell-type annotations. Accurate cell-type annotation is essential for examination of cell-cell interactions and other downstream analyses.

Furthermore, we performed a histological evaluation of the phenotype patterns on each platform identified by pathologists, skilled in immune profiling to reveal any issues that may have been hidden in the large amount of data produced by these platforms. For this assessment, we leveraged H&E staining and mIF with serial sections of samples. The cellular and architectural patterns of tumor cells and nonmalignant cells, including immune cells, are very well known by pathologists and immunologists [19]. We performed this evaluation to validate the data obtained with these platforms, understand the limitations of the outputs of ST platforms, and, most importantly, set the basis for a workflow that can integrate bioinformatics and immune and molecular information with histological categorization of tissues for immune-oncology research.

Although an H&E stain’s ability to show the difference between a CD4 positive and a CD8 positive cell is null, its power to distinguish cell types that have been staples of histology for centuries can be easily examined to increase or gain some confidence in the cell type annotations produced to match the expected cell types in every experimental run. Also, our assessment of the phenotype patterns on ST data was supported by reviewing key biomarkers on mIF images of serial sections of samples to assist pathological evaluation. The ST platforms differed in the number of genes and probes per gene and in gene selection itself, resulting in differences in their ability to produce a dataset that yielded consistent results of their final annotations when compared with each other. Xenium-MM had the highest F1-scores for every phenotype interrogated, whereas MERFISH had the lowest. However, only Xenium allowed us to obtain an H&E staining in the same section used for the assay at the time of our experimental run, thereby facilitating histological evaluation of morphological features and integration of them with gene expression data. These simplified cell type annotations can be challenging to assess without immunology and pathology experience and depending on the experimental design and populations of interest. We recommend visual inspection of the cell in a simplified annotation that encompasses histologically evaluable cell types to evaluate the results of ST data that do not stray from gold standards. This approach can be easily performed on all ST platforms, as they all have the basic lineage gene markers required to perform simplified annotations.

This work has some potential limitations, it was conducted in samples from a single institution, lacks the inclusion of a broader number of tumor types, whole tissue sections, or samples with special tissue processing protocols such as decalcification, these are variables that strongly affect assay performance of any tissue-based assays including single cell ST, further studies considering these special pre-analytical variables are warranted.

## Methods

### Panel design

A predesigned CosMx 1,000-plex panel, MERFISH 500-plex immuno-oncology panel, and Xenium 289-plex human lung panel were selected for this study. Fifty custom genes were added to the Xenium panel: *ARG1*, *B2M*, *CALB2*, *CCL18*, *CCL19*, *CCL2*, *CCL21*, *CCL3*, *CCL4*, *CCL5*, *CCL8*, *CD276*, *CD33*, *CD44*, *CEACAM8*, *CR2*, *CXCL11*, *ENTPD1*, *FAP*, *FCER2*, *HLA-DRB1*, *HSPA6*, *ICOS*, *IDO1*, *IFNG*, *IGHG1*, *IGHG4*, *ITGAX*, *KRT19*, *LAMP3*, *MSLN*, *NCAM1*, *NT5E*, *PTPRC*, *TCF7*, *TGFB1*, *TGFB2*, *TGFB3*, *TIGIT*, *TMEM173*, *TNFRSF9*, *TNFSF4*, *TNFSF9*, *TOX*, *TRDC*, *TREM1*, *VEGFA*, *VSIR*, *VTCN1*, and *ZNF683*.

### Sample preparation

Four TMAs of FFPE tissue from surgically resected tumor samples obtained from patients with lung adenocarcinoma (*n* = 22; 2 TMAs, 53 cores [ICON1 and ICON2]) or pleural mesothelioma (*n* = 22; 2 TMAs, 49 cores [MESO1 and MESO2]) were used. TMA construction was performed using 1-mm-diameter cores from FFPE tumor tissue blocks, with up to three cores per sample. ICON TMA tissues were collected from 2016 to 2018, whereas MESO TMA tissues were collected from 2020 to 2022. All samples were obtained at The University of Texas MD Anderson Cancer Center. This study was approved by the MD Anderson Institutional Review Board and was conducted according to the principles of the Declaration of Helsinki.

TMA cores in the ICON set occupied an area of 17.0 x 13.5 mm in the TMA receiver block, whereas TMA cores in the pleural mesothelioma set occupied an area of 12 x 7 mm.

### Tissue preparation for ST assays

FFPE TMA blocks were sectioned and prepared for each assay according to the ST platform manufacturers’ instructions as described briefly below.

#### CosMx

CosMx uses Superfrost Plus micro slides for placement of tissue. NanoString guidelines required that tissue be placed in the center of a slide in a 20 x-15 mm area, avoiding excess paraffin or tissue on the slide area outside of the scanner area to prevent processing of tissue in a fiducial area [20]. ICON TMAs were scored to select the area of interest, and two TMA rows were excluded for further analysis owing to scanning-area constraints. Pleural mesothelioma TMA blocks were only scored to avoid excess paraffin outside of the scanning area; no cores were excluded. After scoring, sections were cut at 4-5 m and placed in nuclease-free water, excess tissue was removed using a brush, and the selected tissue was aligned at the center of the slide in a 20 x 15-mm area. Slides were then dried at room temperature overnight before storage at 4 °C and then shipping within a week to the NanoString Technology Access Program. Four slides were sent to NanoString for this project.

#### MERFISH

The MERFISH assay is used with special circular slides provided by the manufacturer and requires preparation with an FFPE fiducial premix solution before sectioning in a specific protocol with training required by the provider [21]. Tissue sections were prepared at a thickness of 4-5 m, floated in a nuclease-free water bath, and separated for pickup. The tissue was then picked up, making sure it was within a 12.6 x 15.4-mm area at the center of the slide with the help of a brush. The ICON2 TMA was fitted within the area after excluding one column of tissue from the mounting area. Pleural mesothelioma TMA blocks were only scored to avoid excess paraffin outside the scanning area; no cores were excluded. Also, one TMA (MESO2) was fitted on a single slide. The tissue sections were dried in an oven at 60 °C for 10 min and were then immediately ready for shipping. Slides cushioned in Petri dishes were shipped to Vizgen within 2 weeks of sectioning. The program was restricted to three slides per project.

#### Xenium

Special slides had to be purchased by the manufacturer for tissue placement. These slides had an area of 12 x 24 mm marked by fiducials. After scoring the blocks for the area of interest, each block was sectioned at 4-5 m, floated in a nuclease-free water bath, and separated before picking up the tissue, making sure no part of the tissue or paraffin was touching the fiducials. The ICON2 TMA was scored to select the area of interest (matching the area with the CosMx assay), two TMA rows were excluded for further analysis, and only one TMA (ICON2) fit within the fiducials. Pleural mesothelioma TMA blocks were only scored to avoid excess paraffin outside the scanning area, no cores were excluded, and two TMAs were fitted on a single slide. Next, the slides were incubated at 42 °C for 3 h and dried overnight at room temperature inside a desiccator before shipping [22]. Within 1 week, the slides were sent to the 10x Genomics Catalyst program [23], which was restricted to only two slides per project. Both UM and MM segmentation methods were used with Xenium.

### Data delivery and bioinformatics workflow

The three ST platforms differed in their scanning capabilities and data delivery packages, which required different pipelines and workarounds for downstream analysis. A presentation meeting for the data delivery, consisting of the quality control parameters, representative images of the tissues analyzed, and analysis results with UMAP and phenotype annotations from each assay, was directed by the platform manufacturers’ in-house pathology and data science teams who performed the assays.

#### CosMx

CosMx included whole tissue imaging only for the FOV selection. were manually placed by an operator with a maximum area of 545 by 545 m (at the moment of the experimental run) and selected according to the specific needs and aims of the project. For morphology visualization, mIF with DAPI, PanCk, CD3, and CD45 was performed. The NanoString Spatial Molecular Imaging Technology Access Program had an FOV limit of 25 per slide; however, for this project, the vendor selected a total of 165 FOVs on four slides. For data delivery, NanoString provided the outputs of their in-house pipelines including whole tissue images for FOV selection, high-resolution FL images per FOV, raw channel images of each morphology stain, raw data that included metadata, expression matrix, FOV positions, polygon files that could be used with third-party software, cell composite, cell labels, cell overlay files and Seurat objects, analysis results, napari input files with a custom CosMx plug-in to visualize analysis results, and summary PowerPoint files. Tissue morphology, the captured probes, the FOVs selected for analysis, cell segmentation, and analyzed data from the NanoString in-house pipeline were examined, and morphology scans were exported as OME-TIFF files for use in different digital image analysis suites using scripts within the custom CosMx plug-in in napari.

#### MERFISH

The MERFISH platform has a maximum scan capture area of 1 cm^2^, which must be selected by the user. For data delivery, Vizgen provided the outputs of their in-house pipelines, including cellpose [24] cell-segmentation results in parquet format with selected regions for data analysis using the best morphology stains to generate cell-by-gene and transcript-by-location matrices; low-resolution whole slide images and high-resolution mIF images, consisting of the DAPI nuclear stain and a cocktail of morphology markers labeled as PolyT, Cellbound1, Cellbound2, and Cellbound3; raw images of samples in OME-TIFF file format; data analysis results; and MERFISH input files to explore data using the MERSCOPE Visualizer desktop application.

#### Xenium

The Xenium in situ platform provided full tissue imaging within a scanning area of 12 by 24 mm. For the Xenium-UM assay, slides were stained with only a nuclear stain (DAPI); Later, cell segmentation was performed using an expansion algorithm. For the Xenium-MM assay, a combination of nuclear (DAPI), “Boundary” (ATP1A1, E-cadherin, and CD45), and “Interior” (18S ribosomal RNA) stain cocktails and nuclear expansion up to 5 m assisted the cell-segmentation algorithm. For data delivery, the 10x Genomics Catalyst program provided superimposed H&E-stained images of the TMAs and the raw system outputs, which included the transcript locations without any filtering, as well as segmentation files and analyzed data using their own pipelines to visualize data in Xenium Explorer, which allowed for visualization of the transcripts, and the morphological staining, which included stains for DAPI and a cocktail of visualization markers.

### Data analysis workflow

#### Cell segmentation

Cell-segmentation results for CosMx (enhanced cellpose algorithm [24]), MERFISH (cellpose cell segmentation), and Xenium-MM (custom deep learning models [25]) were provided by the manufacturers, and their outputs were used in the data analysis. However, the cell-segmentation algorithm was rerun for Xenium-UM data with modified parameters using the 10x Genomics Xenium Ranger pipeline (version 1.7.0.2) to eliminate larger cells (-- expansion-distance=7, --dapi-filter=75).

#### Preprocessing, clustering, and annotation

Quality control and data filtering were performed for each ST dataset based on criteria suggested by the ST platform manufacturers. For CosMx, cells with areas more than five times the geometric mean of the cell area, fewer than 30 transcript counts, and FOVs with fewer than an average of 80 transcript counts per cell were filtered. For MERFISH, Xenium-UM, and Xenium-MM, cells with fewer than 10 transcript counts per cell were filtered. These filtering steps eased cell-type annotation while generating gaps between cells.

#### Normalization, dimensional reduction, and cell-type annotation

Standard log-normalization was performed for each dataset using the NormalizeData function in Seurat (version 5) [26]. The dimensionality of each dataset was reduced using the RunPCA function with default parameters and including all targeted genes. TMAs containing samples of the same cancer type were integrated with harmony [27]. The dimensionality was further reduced to two dimensions using the RunUMAP function, and cells were clustered using the FindNeighbors function with the Louvain algorithm and the FindClusters function at 0.8 resolution.

Cells in the ICON TMAs were annotated manually by identifying differentially expressed genes in each cluster using the FindAllMarkers function (Wilcoxon test implemented using the Presto package [28]). The following markers were used to annotate the clusters: *EPCAM* and *PIGR* for tumor cells; *CD19*, *MS4A1* and *BANK1* for B cells; *MZB1* and *JCHAIN* for plasma cells; *CD3E* for T cells, *CD68* and *CD163* for macrophages when the gene expression information was available for the ST platform. Cells were annotated using the label transfer method including the FindTransferAnchors and TransferData functions in Seurat with default parameters from scRNA-seq data of pleural mesothelioma data for the MESO TMAs. Results were evaluated by performing differentially expressed genes in the predicted cell types.

### Bulk RNA-seq and DSP WTA concordance analysis

ST data were correlated with bulk RNA-seq and DSP WTA data of same cohort using Pearson correlation based on shared genes in ST assays.

### Pathological evaluation of cell segmentation and phenotyping

After performing cell-phenotyping annotations for all of the assays, each dataset was organized and compared using a "best possible result" approach to evaluating morphology across the ST platforms. Specifically, the data for each platform were compared by three independent pathologists’ agnostic to all assay information using the ICON2 TMA, which consisted of cores obtained exclusively from patients with non-small cell lung cancer. Cores without tumor tissue were excluded. The pathologists used a simplified annotation method that grouped several of the clusters of the following cell types into morphologically valid features and assigned them colors: tumor cells independent of cluster (cyan), T cells (red), plasma cells (purple), macrophages (orange), fibroblasts and smooth muscle cells (yellow), endothelial cells (green), and other cells (gray or maroon). For CosMx, the cell type annotation was placed next to the H&E stain that was closest to the section of ST data and mIF imaging of the GeoMX WTA assay stained for SYTO13 (DNA), panCK (green, antibody, dilution, fluorophore/channel), and CD3E (red, antibody, dilution, fluorophore/channel) in a composite was performed. For MERFISH, a similar arrangement was used, but the closest H&E-stained section to ST data was a Xenium H&E for the cores where it was available; otherwise, the image came from the next section nearest to ST data. For Xenium, only H&E staining was used for scoring the comparison.

Accuracy scores of 1-5 (1 = best, 5 = worst) were assigned by each pathologist in evaluating the accuracy of the cell-segmentation mask produced by each assay and previously described morphological features. The pathologists were blinded to the diagnosis, tissue core number and position, and each other’s scores during the evaluation. The pathologists’ scores for each ST platform were averaged, with no significant discrepancies.

### Calculation of FDRs and F1-scores

FDR values were calculated for each ST platform using negative control and blank probes individually and the following formula [14–16]:

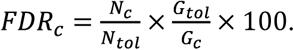

In this formula, *N_tol_* represents the total read count, and *G_tol_* denotes the total number of probes. For assays with negative control probes, *Nc* and *Gc* refer to the total negative control read counts and number of negative control probes, respectively. Similarly, for assays with blank probes, *Nc* and *Gc* corresponded to the total blank read counts and number of blank probes, respectively. F1-scores were calculated for cell segmentation and each cell type, using the average scores of the pathologists as precision scores and the cell-type annotation results as recall with the following formula [18]:

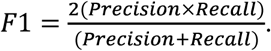

The average scores of the pathologists were normalized by setting precision calls to 1, and the cell-type annotation scores (recalls) were calculated by taking the reciprocal of the normalized average scores. These values were subsequently used to compute F1-scores.

## Acknowledgements

This study was supported by Lung SPORE grant P50 CA070907 from the NIH, the Aileen M. Dillon & Lee M. Bourg Mesothelioma Fund, the Re & RM Kennedy Lung Cancer Fund, the Fleming Endowed Fund, the MD Anderson Rare Tumor Initiative, the MD Anderson Translational Molecular Pathology-Immunoprofiling Lab (TMP-IL) Moon Shots Platform, NCI grant U24CA224285 (to the MD Anderson Cancer Center CIMAC) and the NIH/NCI under award number P30CA016672 and used the Research Histology Core Lab and the Advanced Technology Genomics Core. We thank the MD Anderson Department of Thoracic and Cardiovascular Surgery nurses and staff for their support of this study. The authors acknowledge Donald Norwood, members of the Editing Services, Research Medical Library for editing this manuscript. Fig. 1 was created in BioRender. Ozirmak lermi, N. (2025) https://BioRender.com/y68w242.

## Author contributions

L.M.S.S., C.H., K.C., N.O.L., and M.M.A. conceptualized and designed the experiments. S.H., A.S., K.K., W.L., S.B., M.J., L.M.S.S, RTI Team, J.D., J.Z., B.S., T.C., A.T., M.A., R.Z., D.G., M.G.R., X.T., and M.M.A. prepared and organized data. A.S., S.H., M.M.A., I.L., L.H., L.M.S.S, K.T., RTI Team, J.D., J.Z., B.S., T.C., A.T., M.A., R.Z., D.G., G.R., and X.T. generated data. N.O.L. and K.C. performed bioinformatic data analyses. I.L., L.H., A.S., S.H., L.M.S.S., and M.M.A. performed pathology data analyses. N.O.L. and M.M.A. prepared figures and legends and wrote the manuscript. K.T., B.S., I.W., C.H., L.M.S.S. administered the project. C.H., L.M.S.S, I.W., B.S., acquired funding. C.H., K.C., L.M.S.S supervised the project. All authors reviewed, read, and agreed to publish the manuscript. N.O.L. and M.M.A. contributed equally and all reserve the right to list themselves first on their curricula vitae.

---

**Extended Data Fig. 1.**
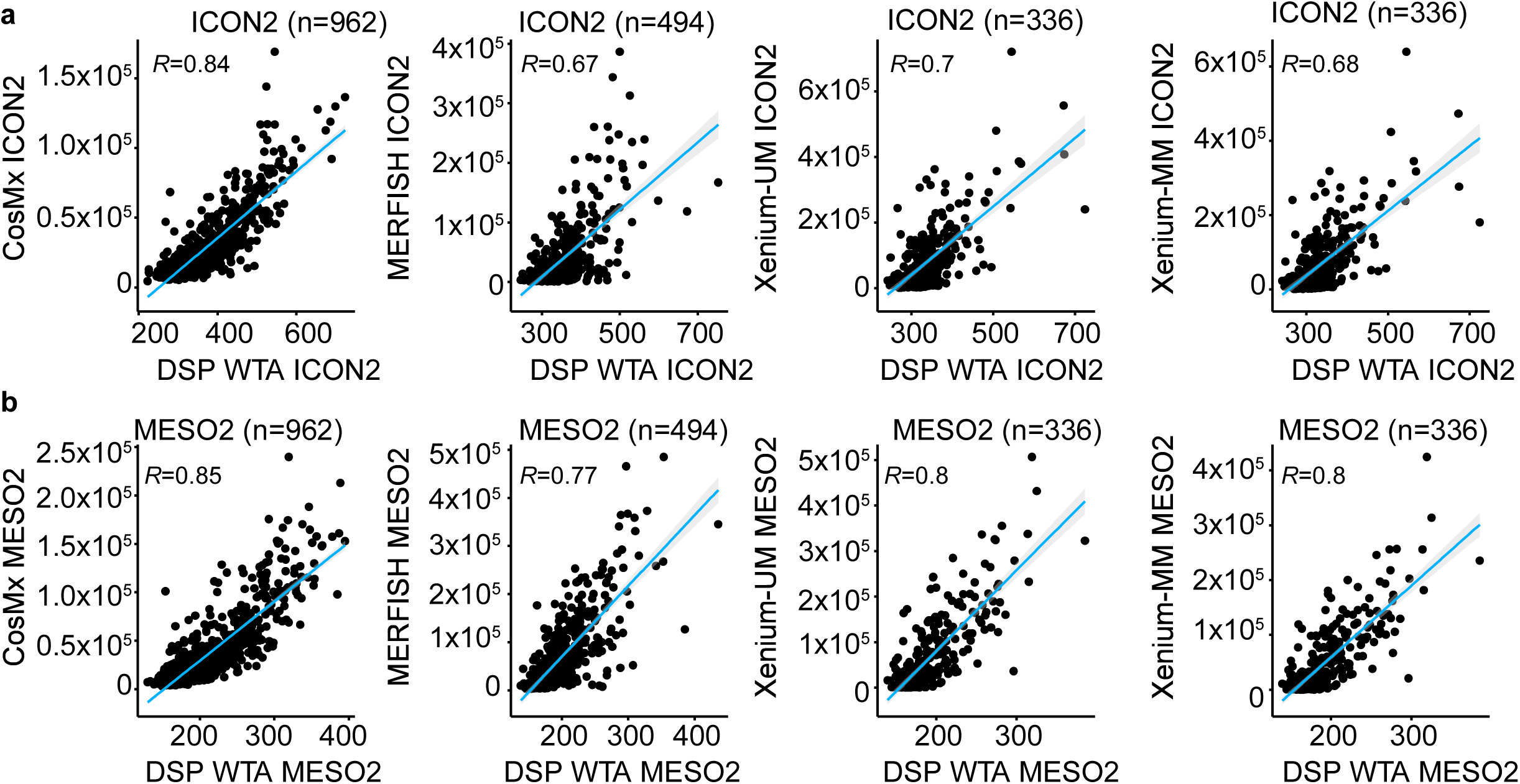
Concordance of ST platforms with DSP WTA. Scatter plots of the average expression of overlapping genes for the ST platforms and DSP WTA with the same cohorts of ICON2 **a,** and MESO2 **b,** TMAs. The blue lines represent linear regression. The Pearson correlation coefficient (*R*) is provided in the top left corner of each plot.

**Extended Data Fig. 2.**
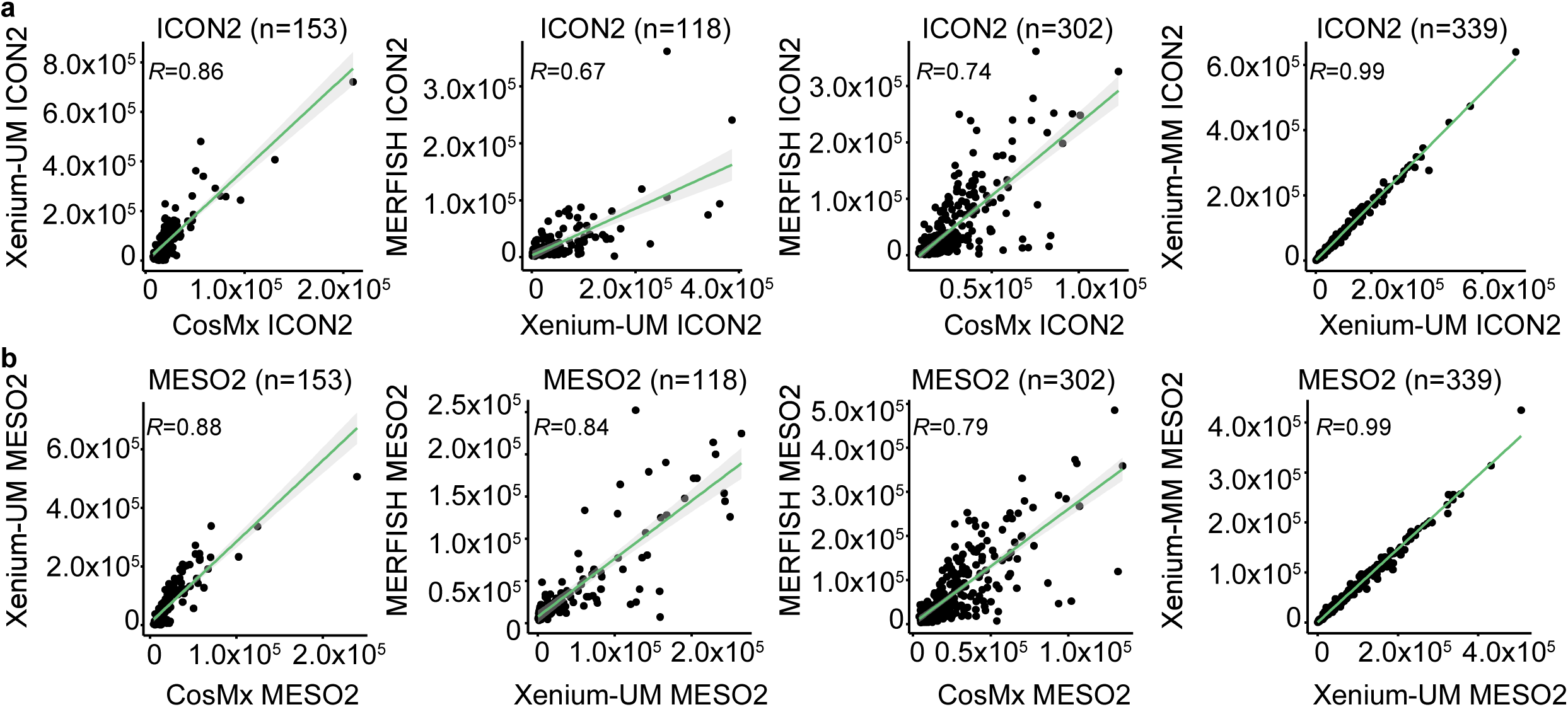
Comparison of gene count detection among the different ST platforms. Scatter plots of the average expression of overlapping genes for the different ST platforms with matched cohorts of ICON2 **a,** and MESO2 **b,** TMAs. The green lines represent linear regression. The Pearson correlation coefficient (*R*) is provided in the top left corner of each plot.

**Extended Data Fig. 3.**
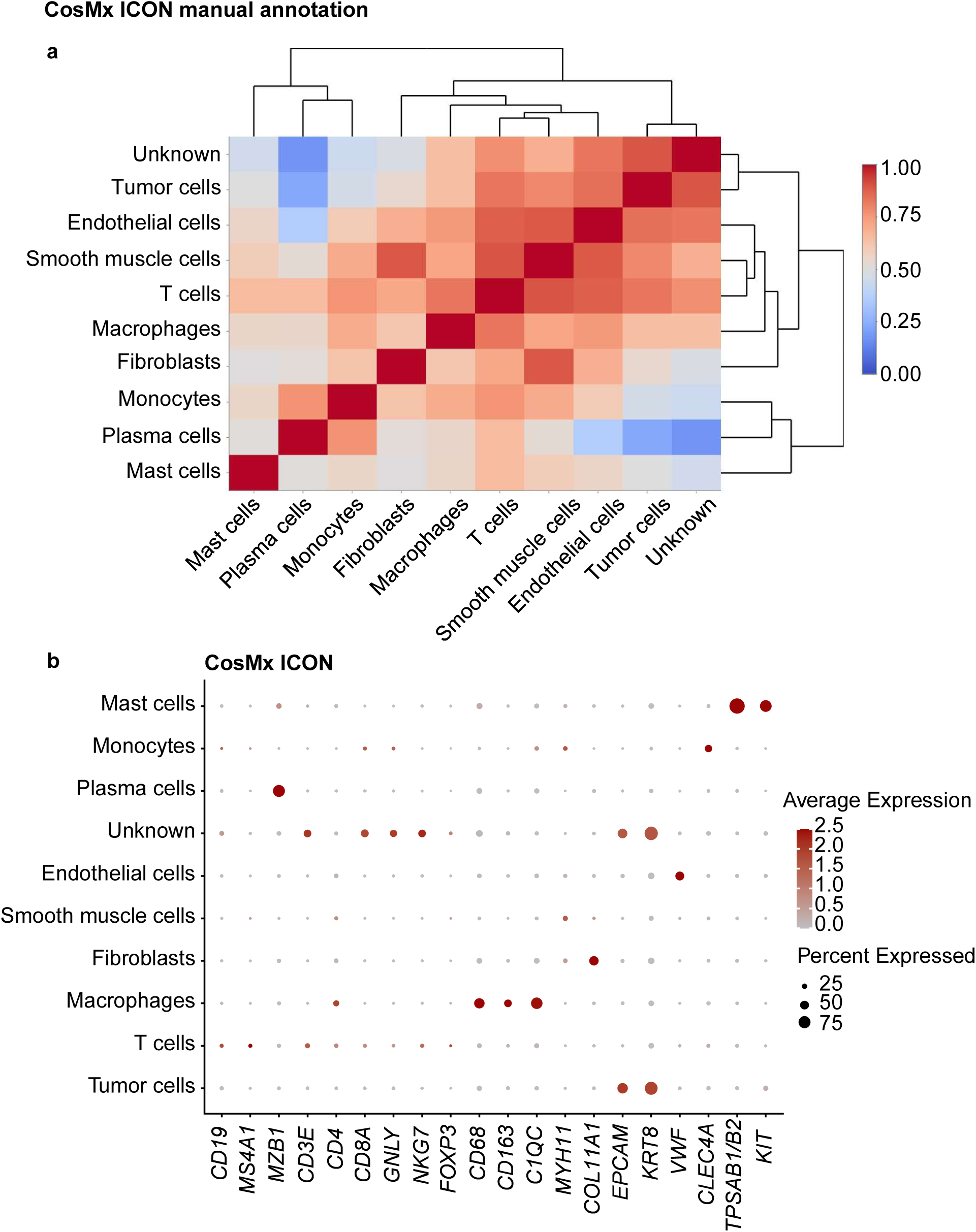
Cell-type annotation for the ICON TMAs on the CosMx platform using a manual cluster annotation approach. **a,** Correlation matrix of the assigned cell types in the ICON TMAs showing high correlation of multiple cell types. Clusters were manually labeled with the corresponding cell types based on the most highly expressed genes in the clusters. **b,** Dot plot of the expression of selected cell marker genes in the assigned cell types in the ICON TMAs.

**Extended Data Fig. 4.**
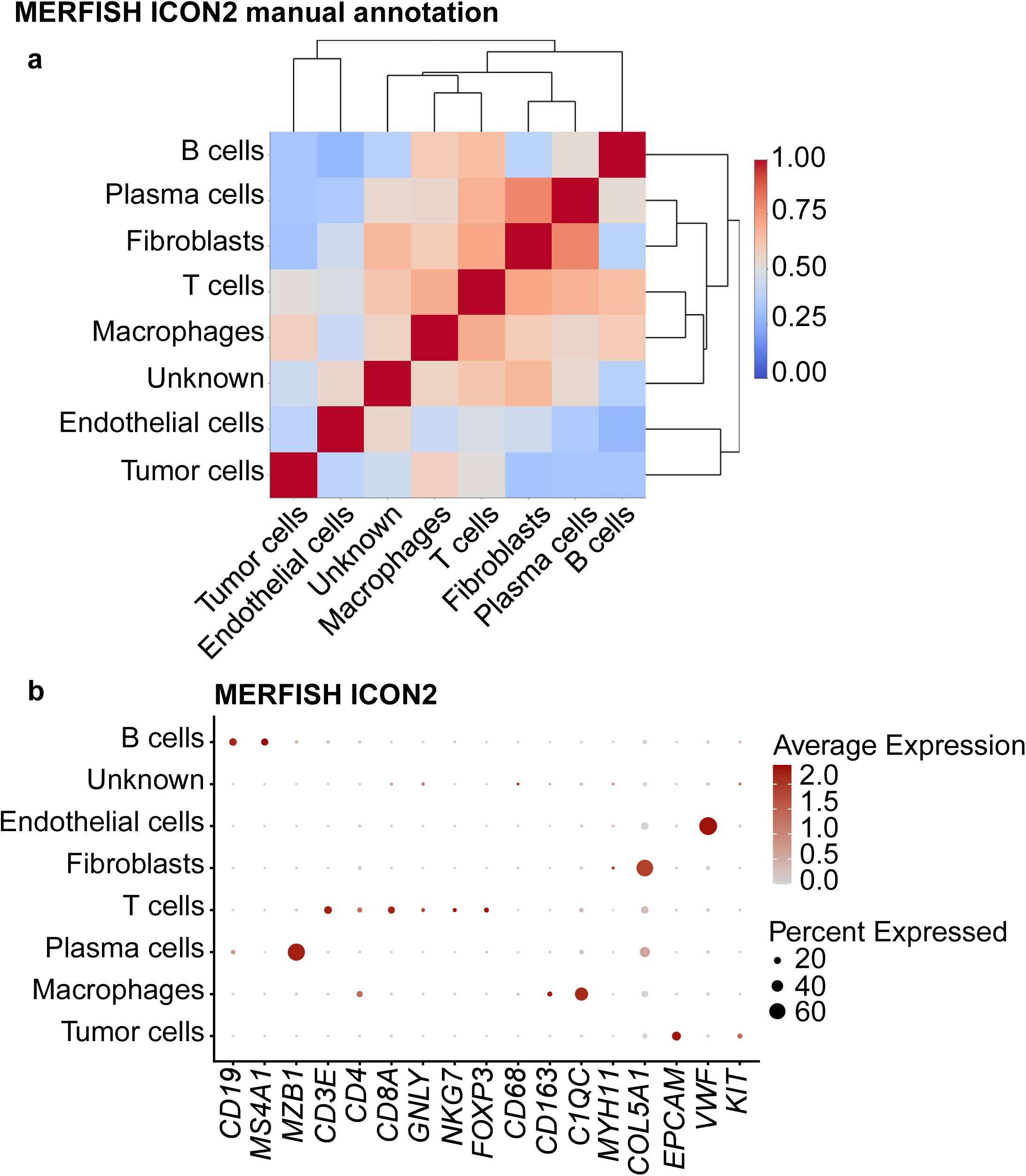
Cell-type annotation for the ICON2 TMA on the MERFISH platform using a manual cluster annotation approach. **a,** Correlation matrix of the assigned cell types in the ICON2 TMA showing high correlation of multiple cell types. Clusters were manually labeled with the corresponding cell types based on the most highly expressed genes in the clusters. **b,** Dot plot of the expression of selected cell marker genes in the assigned cell types in the ICON2 TMA.

**Extended Data Fig. 5.**
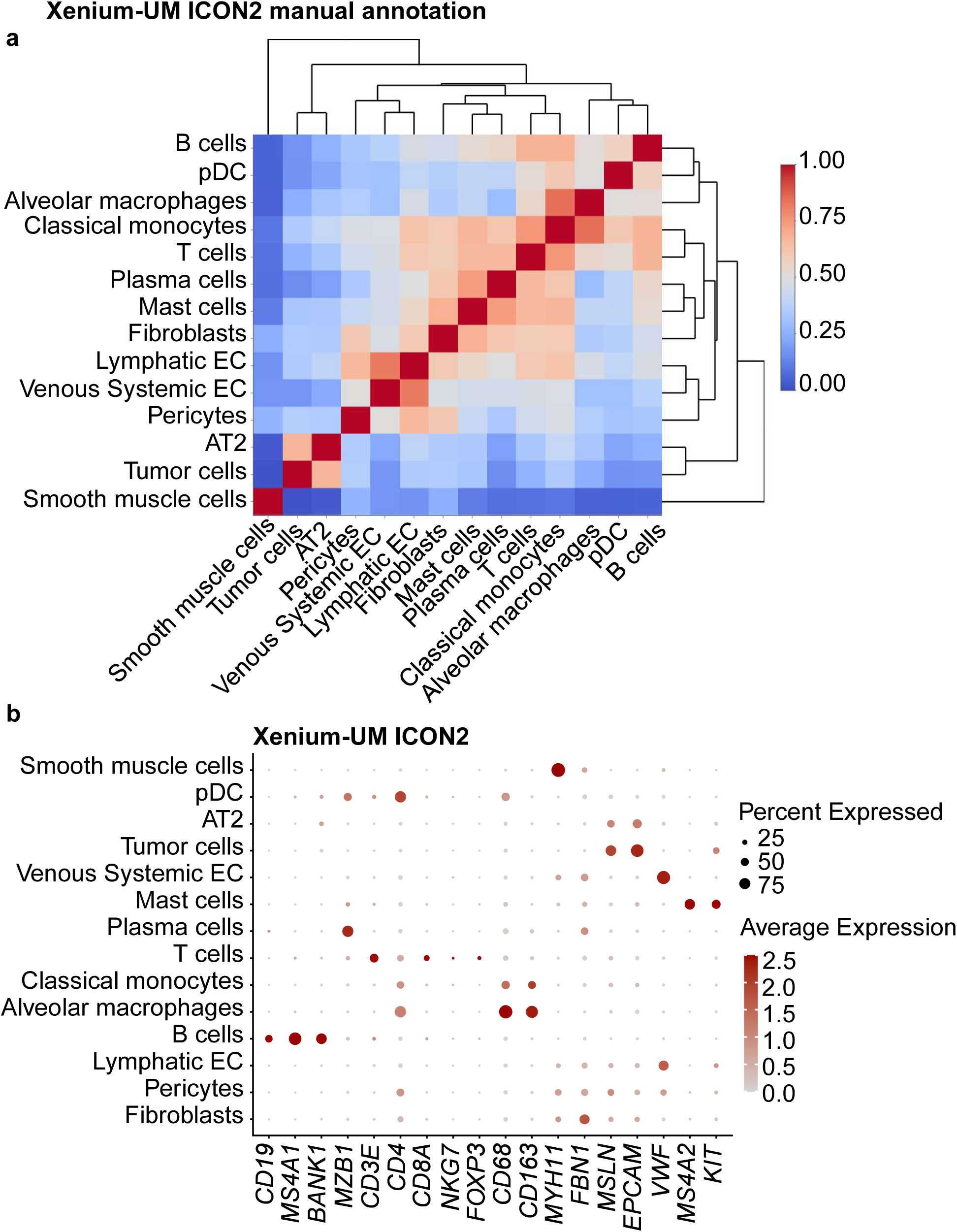
Cell-type annotation for the ICON2 TMA on the Xenium-UM platform using a manual cluster annotation approach. **a,** Correlation matrix of the assigned cell types in the Xenium-UM ICON2 TMA showing distinct separation of multiple cell types. Clusters were manually labeled with the corresponding cell types based on the most highly expressed genes in the clusters. **b,** Dot plot of the expression of selected cell marker genes in the
assigned cell types in the ICON2 TMA.

**Extended Data Fig. 6.**
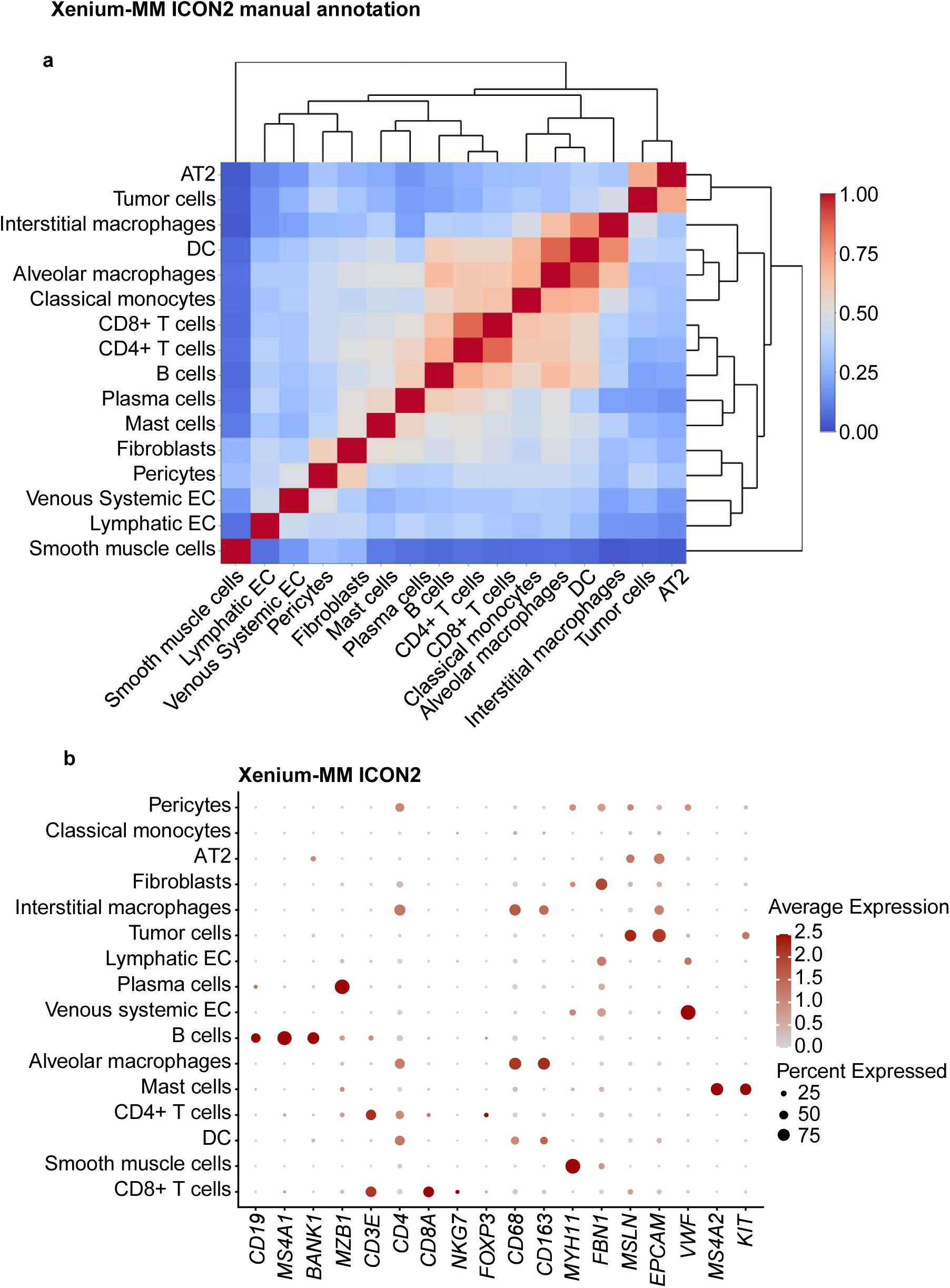
Cell-type annotation for the ICON2 TMA on the Xenium-MM platform using a manual cluster annotation approach. **a,** Correlation matrix of the assigned cell types in the Xenium-MM ICON2 TMA showing distinct separation of multiple cell types. Clusters were manually labeled with corresponding cell types based on the most highly expressed genes in the clusters. **b,** Dot plot of the expression of selected cell marker genes in the
assigned cell types in the ICON2 TMA.

**Extended Data Fig. 7.**
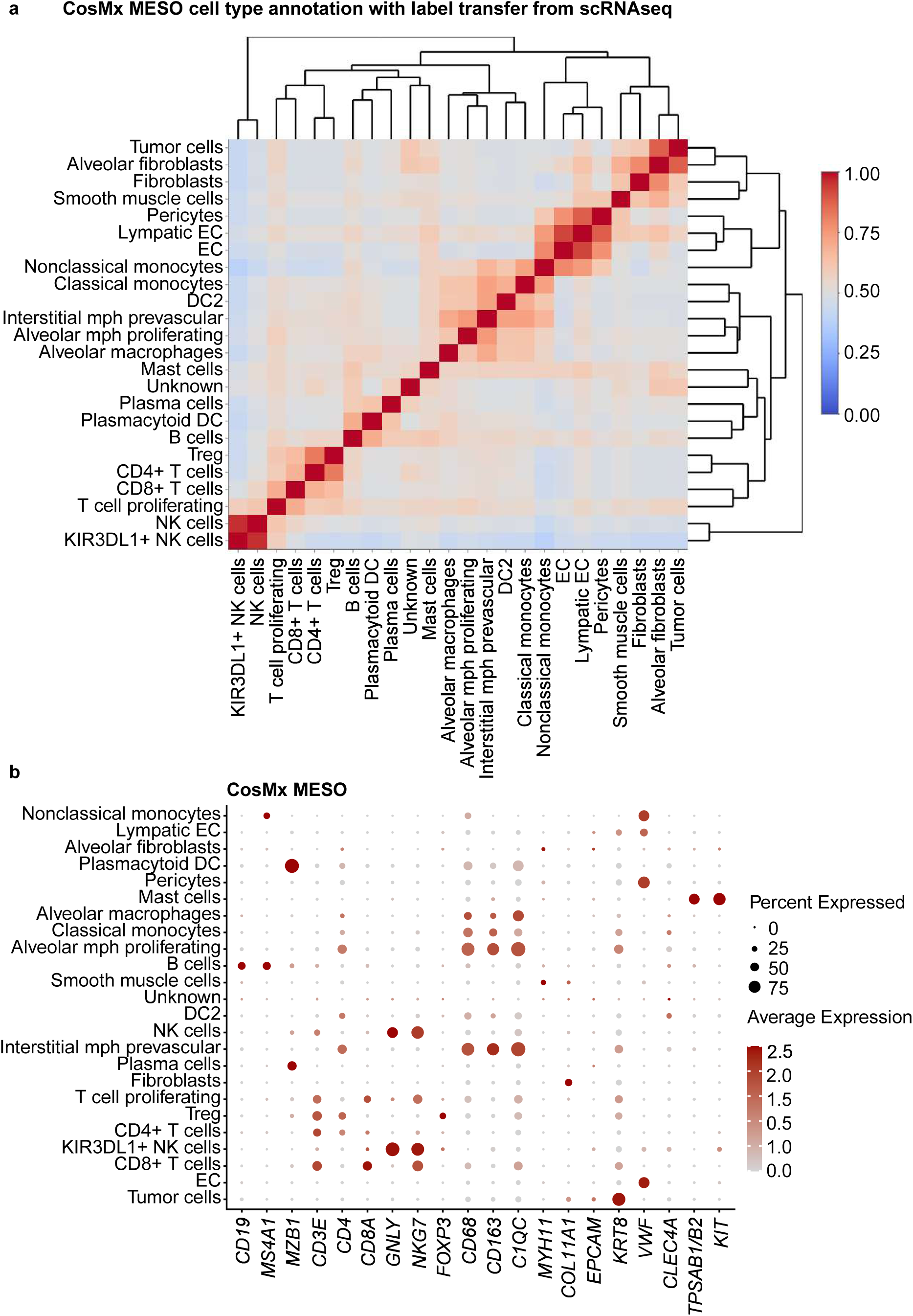
Cell-type annotation for the MESO TMAs on the CosMx platform using a transfer label from scRNA-seq. **a,** Correlation matrix of the assigned cell types in the CosMx MESO TMAs showing distinct separation of multiple cell types. Cell types were labeled using transferring label approach from scRNA-seq data. **b,** Dot plot of the expression levels for selected cell marker genes in the assigned cell types.

**Extended Data Fig. 8.**
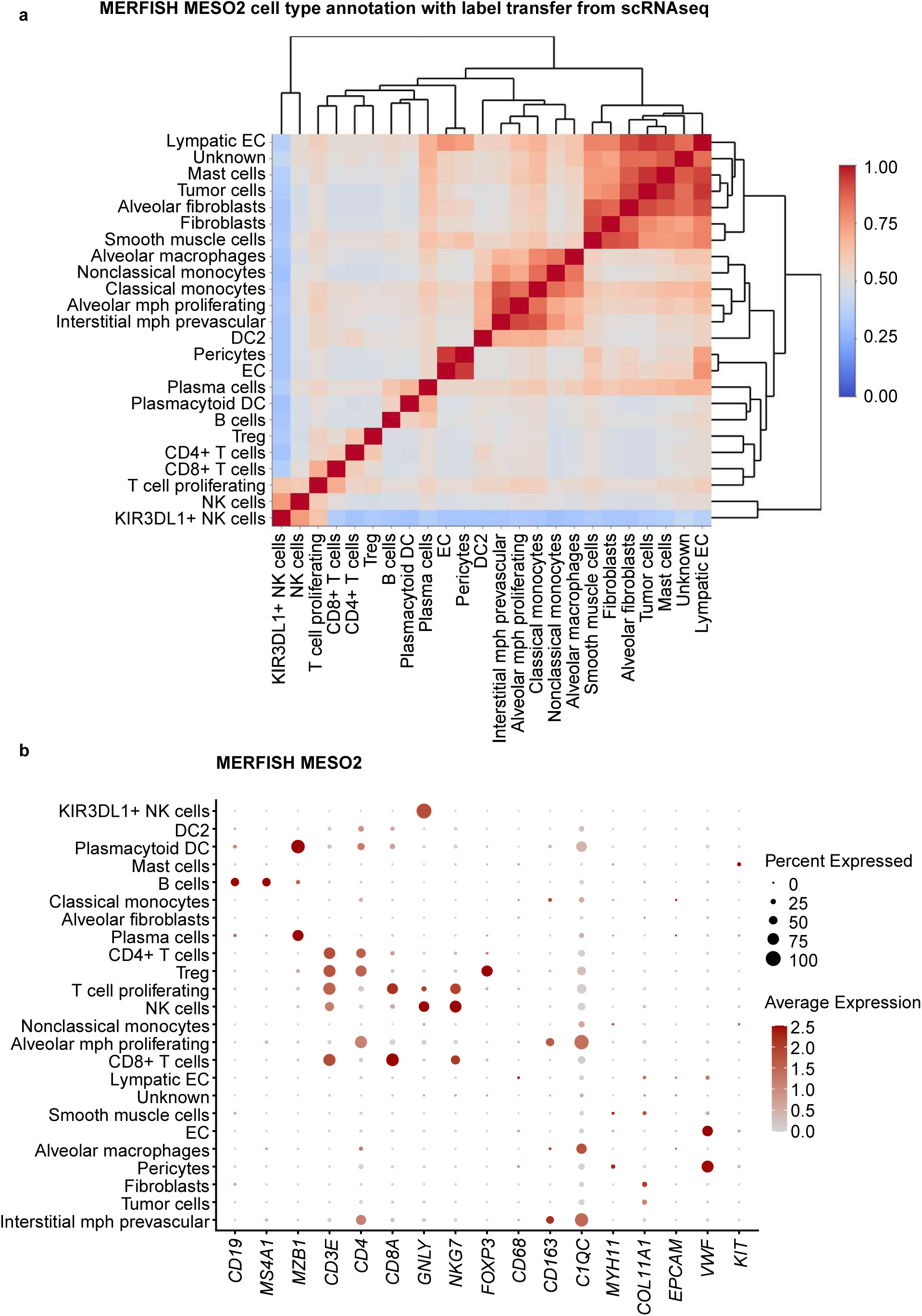
Cell-type annotation for the MESO2 TMA on the MERFISH platform using transferring label approach from scRNA-seq data. **a,** Correlation matrix of the assigned cell types in the MERFISH MESO2 TMA showing distinct separation of multiple cell types. Cell types were labeled using a transferring label approach from scRNA-seq data. **b,** Dot plot of the expression levels for selected cell marker genes in the assigned cell types.

**Extended Data Fig. 9.**
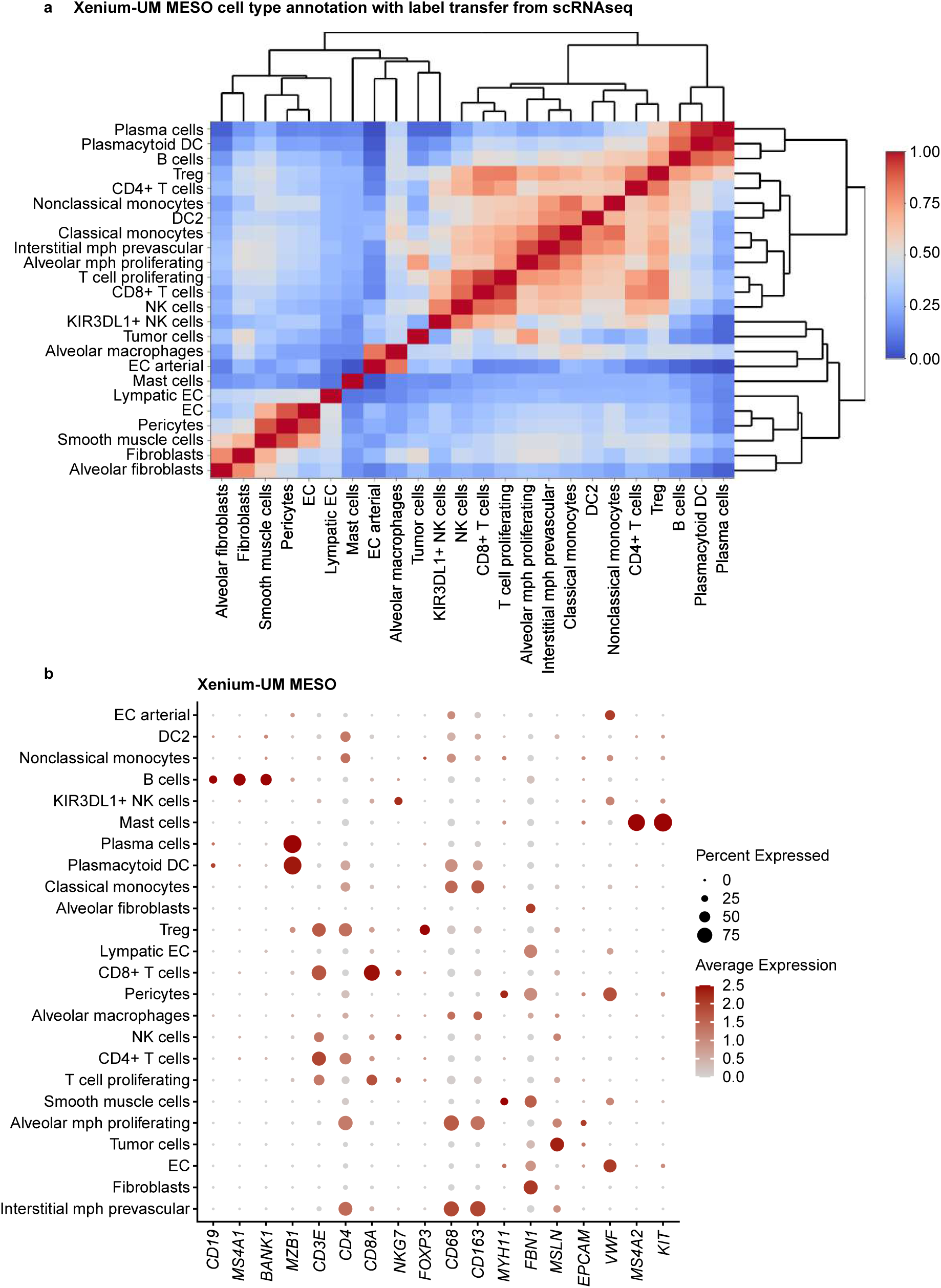
Cell-type annotation for the MESO TMAs on the Xenium-UM platform using a transfer label from scRNA-seq. **a,** Correlation matrix of the assigned cell types in Xenium-UM MESO TMAs showing distinct separation of multiple cell types. Cell types were labeled using a transferring label approach from scRNA-seq data. **b,** Dot plot of the expression levels for selected cell marker genes in the assigned cell types.

**Extended Data Fig. 10.**
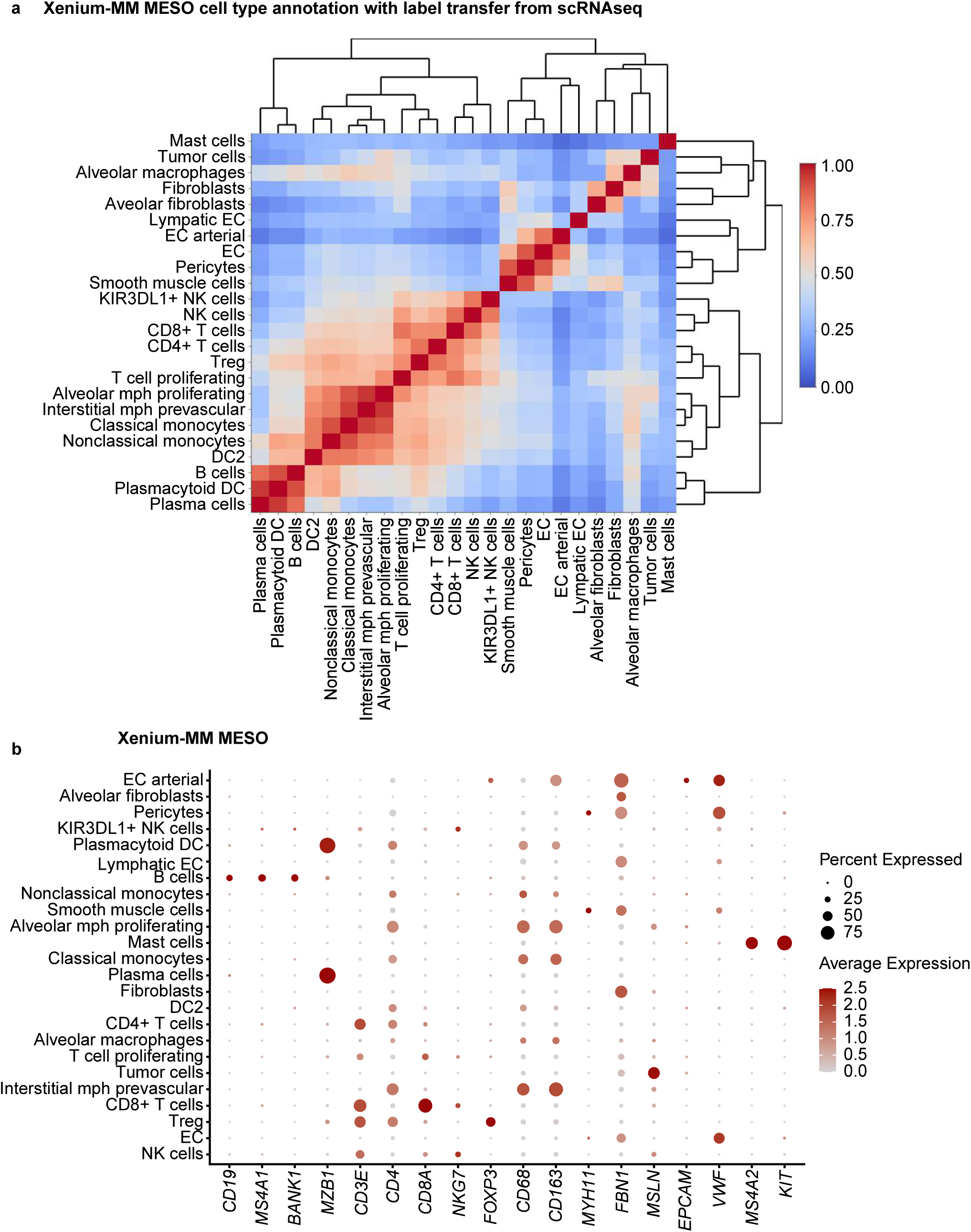
Cell-type annotation for the MESO TMAs on the Xenium-MM platform using a transfer label from scRNA-seq. **a,** Correlation matrix of the assigned cell types in Xenium-MM MESO TMAs showing distinct separation of multiple cell types. Cell types were labeled using a transferring label approach from scRNA-seq data. **b,** Dot plot of the expression levels for selected cell marker genes in the assigned cell types.

