## Supplementary Figure 2 for "Comparison of imaging-based single-cell resolution spatial transcriptomics profiling platforms using formalin-fixed, paraffin-embedded tumor samples"

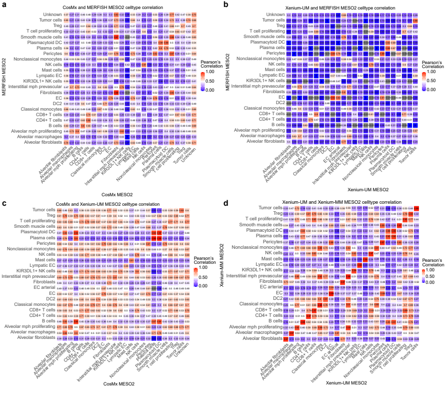


**Supplementary Figure 2: Correlation of annotated cell types between ST platforms in MESO2.** Heatmaps show correlation of annotated cell types between **(a)** CosMx and MERFISH, **(b)** Xenium-UM and MERFISH, **(c)** CosMx and Xenium-UM and **(d)** Xenium-UM and Xenium-MM of MESO2 based on shared genes. Red and blue colors represent high and low Pearson's correlation between celltypes, repectively, Correlation coefficients are written on heatmap cells.
