## Supplementary Figure 3 for "Comparison of imaging-based single-cell resolution spatial transcriptomics profiling platforms using formalin-fixed, paraffin-embedded tumor samples"

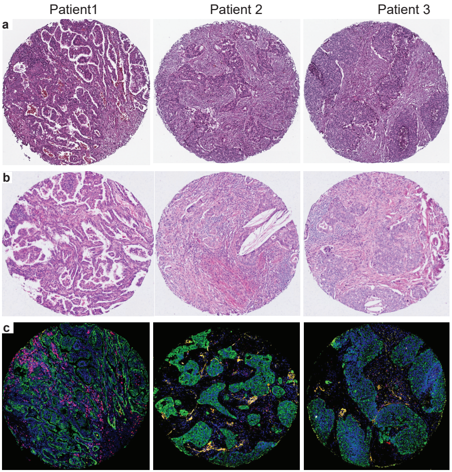


**Supplementary Figure 3: Strategy of pathologists' evaluation on ICON2.** Microphotographs showing H&E **(a)** , reference H&E, **(b)** Xenium-MM H&E staining slides, **(c)** multiplex immunofluorescence stained with CK (green), CD3 (red) and CD68 (yellow) performed in a serial sections. We evaluated the pattern of expression of endothelial cells, fibroblasts, macrophages, plasma cells, T cells and epithelial cells, and compared them with the patterns of the phenotypes obtained for each platform in ICON2.
